# Spatial and Multi-Omics Analysis of Human Breast Cancer Reveals the Spatiotemporal Dynamics of Basal Layer Disruption

**DOI:** 10.64898/2026.08.10.744069

**Authors:** Siwei Ju, Zhanzhi Li, Lidan Jin, Yongxia Chen, Jun Shen, Mengqing Liu, Yi Lu, Qina He, Xixi Lin, Dazhi Chen, Jiahang Zhang, Menghao Zhou, Qingqing Fang, Qingqing Hu, Yuanhang Xia, Qingna Meng, Jiaheng Lang, Lili Zhi, Xia Yang, Yongtao He, Yihong Wen, Xuanjing Ye, Jichun Zhou, Linbo Wang, Feiyang Ji

**Author notes:** Correspondence (F.J.), (L.W.), (J.Z.). These authors contributed equally (Siwei Ju, Zhanzhi Li, Lidan Jin, Yongxia Chen, Jun Shen, Mengqing Liu).

## Abstract

When breast cancer invasion begins and how tumor cells breach the basal barrier remain poorly defined. We profiled normal mammary ducts, ductal hyperplasia (DH), ductal carcinoma in situ (DCIS) and invasive ductal carcinoma using spatial transcriptomics, spatial proteomics and five bulk-omics layers, alongside an independent longitudinal lesion cohort. Cross-sectionally, basal/myoepithelial continuity declined most between DH and DCIS, accompanied by extracellular-matrix remodeling and altered fibroblast- and macrophage-associated signaling. EGFR-positive luminal progenitor-like cells were enriched at manually annotated basal discontinuities and were molecularly distinct, nominating a candidate leader-like population without establishing causality. In the longitudinal cohort, expression of GABRG3, TAGLN, MLPH and AZGP1 in initially benign lesions was associated with subsequent ipsilateral malignancy. These findings support a model in which progression-relevant breast tissue remodeling may begin at the DH stage and nominate cellular states and candidate biomarkers for prospective validation in breast cancer risk stratification among patients with DH.

---

Breast cancer is one of the most common malignancies and a leading cause of cancer-related mortality among women worldwide^1,2^. With the widespread implementation of breast cancer screening and advances in imaging technologies, an increasing number of patients are diagnosed with proliferative breast lesions, including ductal hyperplasia (DH). Although epidemiological studies have shown that certain DH lesions are associated with an elevated risk of subsequent breast cancer development^3–5^, reliable biomarkers and risk assessment strategies capable of distinguishing lesions that remain indolent from those that progress to ductal carcinoma in situ (DCIS) or invasive ductal carcinoma (IDC) are still lacking. Identifying high-risk DH lesions in which malignant transformation has already been initiated and establishing precise risk stratification strategies therefore remain major challenges in the clinical management of breast disease.

Breast cancer progression is generally thought to follow a stepwise evolutionary trajectory from normal mammary ducts to DH, DCIS, and ultimately IDC^6^. However, increasing evidence suggests that histological boundaries do not necessarily reflect discrete biological states. DCIS, the principal precursor of IDC, carries a substantial risk of progression to invasive disease if left untreated^7^. Moreover, molecular abnormalities associated with tumorigenesis have been detected in lesions that are still histologically classified as benign^8^. These observations raise the possibility that malignant transformation programs may be initiated before the onset of conventionally defined premalignant stages. However, it remains unclear whether the earliest malignant events emerge during DCIS, DH, or even earlier stages, and which biological processes underlie the markedly different clinical outcomes observed among DH lesions. Defining the temporal origin of malignant transformation therefore represents a fundamental challenge in understanding early breast tumorigenesis and developing robust risk prediction strategies.

A hallmark event during breast cancer invasion is disruption of the basal/myoepithelial barrier. The normal mammary duct consists of an inner luminal epithelial layer and an outer basal/myoepithelial layer. Luminal epithelial cells comprise both luminal progenitor cells, which possess substantial proliferative capacity and lineage plasticity, and mature luminal cells responsible for secretory functions^9^. Together with the basement membrane, the basal/myoepithelial layer forms a continuous bilayered barrier that maintains epithelial polarity, regulates extracellular matrix homeostasis, and suppresses tumor progression^10,11^. Consequently, loss of basal layer integrity has long been regarded as a defining feature of invasive breast cancer^12–14^. Notably, accumulating evidence suggests that invasion-associated programs may arise before histological invasion becomes apparent. Focal disruption of the myoepithelial layer and basement membrane has been reported in non-invasive lesions^15,16^, while studies supporting early dissemination and parallel progression models have further challenged the traditional view that invasion strictly follows a linear evolutionary trajectory^17–19^. Nevertheless, when basal barrier disruption is initiated, which cellular populations drive this process, and how the local microenvironment contributes to malignant transformation remain poorly understood in human tissues.

A major obstacle to addressing these questions is the lack of human model systems that faithfully capture the continuous evolutionary process of breast cancer. Although animal models have provided important insights into breast cancer biology^20,21^, they do not fully recapitulate the architectural complexity and evolutionary dynamics of human breast tissues. Recent advances in spatial multi-omics technologies have enabled the characterization of cellular states and intercellular communication networks at single-cell resolution while preserving tissue architecture, thereby facilitating reconstruction of the spatial ecosystems that govern tumor evolution^22–24^. However, a comprehensive spatial and multi-omics analysis spanning normal mammary ducts, DH, DCIS, and IDC in human tissues has not yet been achieved.

To address this gap, we established two complementary human breast cancer cohorts. First, from more than 400 breast cancer specimens, we identified over 100 samples containing multiple pathological stages within the same breast tissue and assembled a breast cancer evolution cohort spanning normal tissue from malignant patients (MN), DH, DCIS, and IDC. Using a space-for-time design, we compared spatial and molecular features across pathological stages. Second, we established an independent clinical outcome cohort comprising 90 patients initially diagnosed with benign breast lesions who subsequently developed either benign or malignant ipsilateral breast tumors. By integrating single-cell spatial transcriptomics, single-cell spatial proteomics, whole-exome sequencing, transcriptomics, proteomics, metabolomics, and microbiome profiling, we profiled stage-associated spatial and molecular patterns across the cohort. We found that invasion-associated molecular programs emerge during the DH stage rather than at the DCIS or IDC stage. Across the DH and DCIS stages, basal layer disruption became increasingly evident and was accompanied by extensive remodeling of the local microenvironment. We further identified EGFR-positive luminal progenitor-like cells that may act as leader cells driving early breaching of the basal barrier. In addition, we identified spatial molecular features in DH lesions associated with subsequent malignant progression and validated their predictive value in an independent clinical outcome cohort. Together, these findings support a model in which breast cancer progression may be initiated at the DH stage and provide a foundation for risk stratification and precision management of patients with DH.

## Results

### 1 Study design and integrated spatial multi-omics profiling

To systematically characterize the molecular and spatial features underlying breast cancer progression, we established a multi-omics cohort spanning normal, hyperplastic, in situ, and invasive disease stages (Figure 1A). For patients undergoing mastectomy, multiple tissue specimens were collected from different regions of the resected breast and classified by pathological review as MN, DH, DCIS, or IDC. Because pathological stages varied among patients, not all specimens contained the complete sequence of disease progression. Therefore, only patients with at least three pathological stages represented within the same breast were included in the discovery cohort. Following pathological review and stringent quality control, the final cohort comprised 92 tissue samples from 27 mastectomy patients, including 22 MN, 23 DH, 15 DCIS, and 32 IDC samples. In addition, normal mammary duct tissues from 10 fibroadenoma (FA) patients were included as benign controls and designated as normal tissue from benign patients (BN). Each tissue specimen was divided into two portions. One portion was snap-frozen and subjected to bulk multi-omics profiling, including whole-exome sequencing (WES), transcriptomics, proteomics, 16S microbiome profiling, and metabolomics. The remaining portion was formalin-fixed and paraffin-embedded for tissue microarray construction and subsequent single-cell spatial transcriptomic (CosMx) and spatial proteomic (CODEX) analyses. Multi-omics findings were further evaluated using tissue microarrays and multiplex immunohistochemistry (mIHC) in an independent validation cohort.

**Figure 1.**
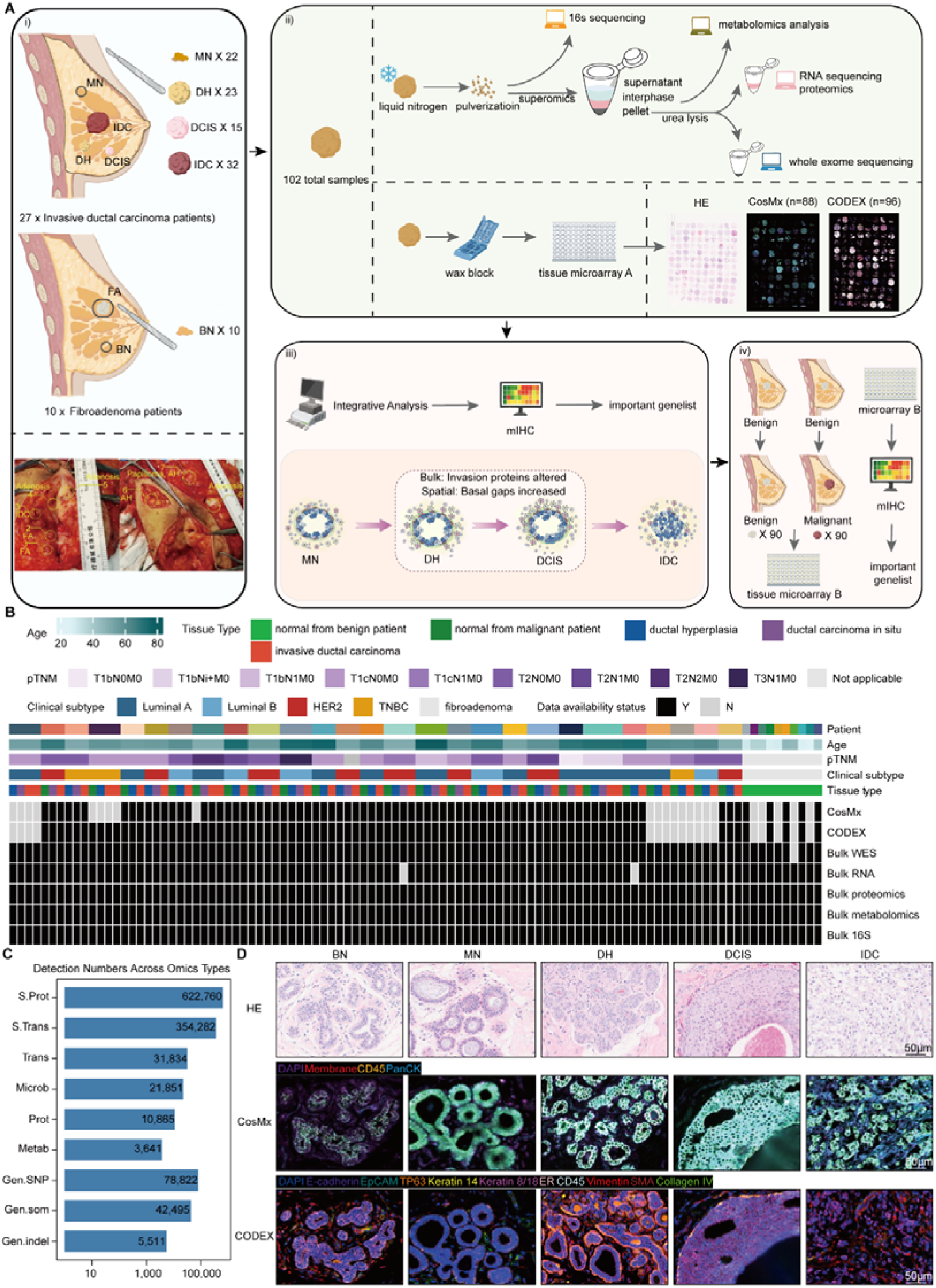
Study design and integrated spatial multi-omics profiling. (A) Schematic illustrating the workflow, including samples collection, data acquisition, integrative analysis and validation of key findings. (B) Heatmap of patient/sample clinical metadata and multi-omics data availability. Columns represent individual tissue samples; rows denote patient ID, age, pTNM stage, clinical subtype, tissue type, and assay coverage for CosMx, CODEX, bulk WES, bulk RNA-seq, bulk proteomics, bulk metabolomics, and bulk 16S rRNA profiling. (C) Number of detected features across different omics modalities, including spatial proteomics, spatial transcriptomics, transcriptomics, microbiome, proteomics, metabolomics, and genomic alterations. (D) Representative H&E staining and matched CosMx and CODEX spatial profiles from serial adjacent tissue sections across breast cancer progression stages. Scale bars, 50 μm. BN, normal from benign patient; MN, normal from malignant patient; DH, intraductal hyperplasia; DCIS, ductal carcinoma in situ; IDC, invasive ductal carcinoma; FA, fibroadenoma; AH, atypical hyperplasia; mIHC, multiplex immunohistochemical; S. Prot, spatial proteomics; S. Trans, spatial transcriptomics; Trans, transcriptomics; Microb, microbiome; Prot, proteomics; Metab, metabolomics; Gen. SNP, genomic single nucleotide polymorphisms; Gen. som, genomic somatic mutations; Gen.indel, genomic indel.

The mean age of breast cancer patients was 59.9 years. Molecular subtypes reflected the distribution observed in real-world clinical populations and included 11 Luminal A, 5 Luminal B, 8 HER2-positive, and 3 triple-negative breast cancer. Most patients presented with stage I-II disease. Notably, six patients contained all four pathological stages, including MN, DH, DCIS, and IDC, within the same breast specimen (Figure 1B). CosMx profiling was performed on 79 samples across 386 fields of view (FOVs), whereas 84 samples underwent CODEX analysis. In parallel, 101 samples were subjected to WES, 100 samples underwent transcriptome sequencing, and 102 samples were analyzed by microbiome, proteomic, and metabolomic profiling (Figure 1B; Table S1). We obtained spatial transcriptomic data from 354,282 cells and spatial proteomic data from 622,760 cells. Bulk multi-omics profiling identified 42,495 somatic mutations, 5,511 insertions/deletions (indels), 78,822 single-nucleotide polymorphisms (SNPs), 31,834 transcripts, 10,865 proteins, 21,851 microbial features, and 3,641 metabolites (Figure 1C).

Representative histological and spatial multi-omics images revealed marked differences in tissue architecture and cellular composition across disease stages (Figure 1D). Integrative analysis of hematoxylin and eosin (H&E) staining, CosMx, and CODEX data captured the dynamic remodeling of tissue structure and cellular spatial organization during progression from MN and DH to DCIS and IDC, providing a high-resolution spatial framework for investigating ecosystem remodeling throughout breast cancer evolution.

### 2 Dynamic changes in spatial cellular composition during breast cancer progression

After excluding cells with abnormal segmentation, we retained 334,025 high-quality cells from the CosMx dataset and 606,467 high-quality cells from the CODEX dataset for downstream analyses (Figure S1). Six major cell populations were identified based on canonical marker genes, including epithelial cells, endothelial/perivascular cells, fibroblasts, myeloid cells, T/B/NK cells, and plasma cells in CosMx (Figures 2A and 2B). Epithelial cells were characterized by high expression of *EPCAM*, *KRT19* and *KRT18*, whereas fibroblasts were enriched for stromal markers such as *COL1A2* and *COL3A1*. Immune cell populations exhibited lineage-specific expression patterns consistent with their cellular identities. For the CODEX dataset, unsupervised Leiden clustering (resolution = 0.5) identified five major cell populations, including epithelial cells, endothelial/perivascular cells, fibroblasts, myeloid cells, and T/B cells (Figures 2C, 2D).

**Figure 2.**
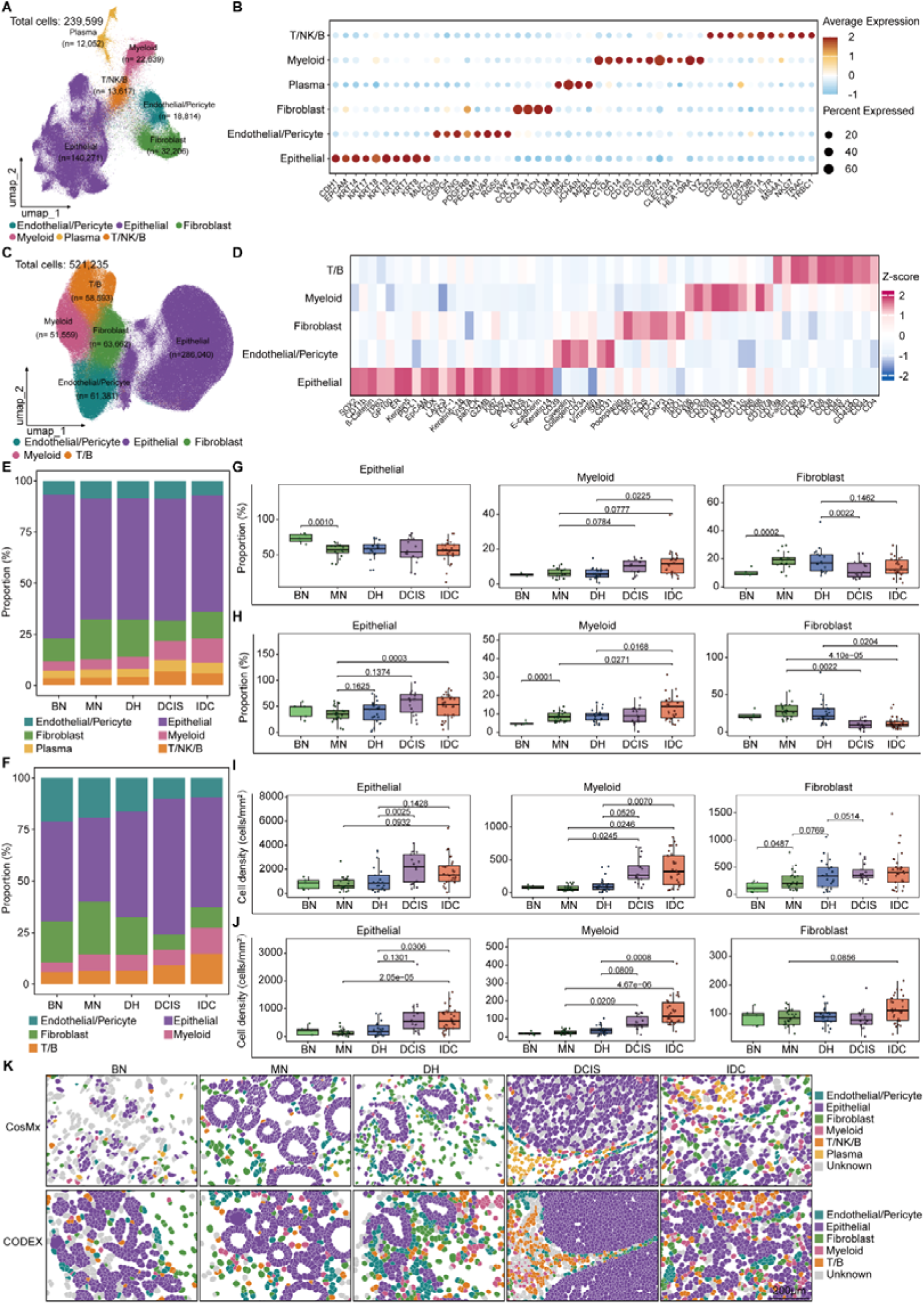
Dynamic changes in spatial cellular composition during breast cancer progression. (A) UMAP projection of 239,599 cells from CosMx spatial transcriptomics, colored by major cell types. (B) Dot plot showing average expression level (color) and percentage of expressing cells (size) of marker genes for each major cell type in CosMx data. (C) UMAP projection of 521,235 cells from CODEX spatial proteomics, colored by major cell types with a consistent color scheme as CosMx. (D) Heatmap of Z-score normalized marker protein expression across major cell types in CODEX data. (E) Stacked bar plot showing the proportion of major cell types across benign normal (BN), malignancy normal (MN), intraductal hyperplasia (DH), ductal carcinoma in situ (DCIS) and invasive ductal carcinoma (IDC) stages in CosMx data. (F) Stacked bar plot showing the proportion of major cell types across BN, MN, DH, DCIS and IDC stages in CODEX data. (G) Box-and-scatter plots depicting the cell proportion of epithelial, myeloid and fibroblast cells across disease stages in CosMx data. (H) Box-and-scatter plots depicting the cell proportion of epithelial, myeloid and fibroblast cells across disease stages in CODEX data. (I) Box-and-scatter plots showing cell density (cells/mm^2^) of epithelial, myeloid and fibroblast cells across disease stages in CosMx data. (J) Box-and-scatter plots showing cell density (cells/mm^2^) of epithelial, myeloid and fibroblast cells across disease stages in CODEX data. (K) Representative spatial distribution maps of major cell types in CosMx (top) and CODEX (bottom) data across all disease stages. Scale bar, 200 μm. Statistical analysis: Seven pairwise stage comparisons were conducted. Independent-sample tests were used for MN vs BN, whereas paired tests were applied to the other six comparisons using only matched patient samples (DH vs MN, DCIS vs DH, IDC vs DCIS, DCIS vs MN, IDC vs MN, IDC vs DH). All tests were performed using the t test for G-J. Comparisons without indicated *P* values were not statistically significant. *P* values are presented to four significant digits; *P <* 0.0001 is indicated in scientific notation. BN, normal from benign patient; MN, normal from malignant patient; DH, intraductal hyperplasia; DCIS, ductal carcinoma in situ; IDC, invasive ductal carcinoma.

We next examined cellular composition across disease stages to characterize large-scale ecosystem remodeling during breast cancer progression (Figures 2E, 2F). Although the proportion of epithelial cells showed only modest changes across stages in the CosMx dataset, both CosMx and CODEX data revealed a marked increase in epithelial cell density that peaked at the DCIS stage. This finding is consistent with extensive intraductal epithelial expansion during in situ tumor growth. Immune cell populations also exhibited progressive accumulation during disease progression. Both the abundance and density of myeloid cells and T/B/NK cells increased from benign to malignant stages, with the most pronounced increase in myeloid cells observed in DCIS and IDC. These findings indicate continuous remodeling of the immune microenvironment during tumor evolution. Endothelial/perivascular cells displayed a stage-dependent non-linear pattern. Their density progressively increased from MN to DCIS but declined in IDC. Both cellular proportion and density were significantly lower in IDC than in DCIS. In contrast, fibroblasts showed a marked reduction in relative abundance at malignant stages, whereas their cellular density remained largely unchanged (Figures 2G–2J and S2A–S2D). These observations suggest that the reduced fibroblast proportion primarily reflects the expansion of epithelial and immune compartments rather than an absolute loss of fibroblasts. Together, these changes are consistent with emerging evidence that enhanced immune infiltration and stromal remodeling jointly shape the evolving tumor ecosystem during breast cancer progression^25,26^. To validate these quantitative findings in situ, we examined the spatial distribution of major cell populations using both CosMx and CODEX (Figure 2K). Spatial organization closely mirrored the quantitative analyses. BN and MN samples maintained relatively well-organized ductal structures, whereas DH, DCIS, and IDC lesions progressively exhibited epithelial expansion, immune cell enrichment, and stromal remodeling. These findings further support dynamic ecosystem reorganization throughout breast cancer progression.

In addition to compositional changes, we investigated cell morphology across disease stages (Figures S2E, S2F; Table S2). In the CosMx dataset, epithelial cells in IDC displayed significantly larger cell areas and increased major-axis length, suggesting cellular enlargement and structural abnormalities associated with invasive progression. These features are consistent with the loss of epithelial polarity and enhanced invasiveness of tumor cells. By contrast, fibroblasts in DCIS exhibited reduced cell area, long axis length, and short axis length compared with those in DH, accompanied by decreased eccentricity. This finding indicates substantial morphological remodeling of fibroblasts during the transition from DH to DCIS.

Finally, we integrated spatial omics findings with bulk transcriptomic and proteomic datasets. Expression of the epithelial marker EPCAM showed minimal changes across disease stages in both bulk and spatial datasets, consistent with the relatively stable proportion of epithelial cells observed at the single-cell level (Figures S2G and S2H). In contrast, individual epithelial markers displayed distinct trajectories, with *KRT8* progressively increasing and *KRT5* progressively decreasing, suggesting dynamic shifts among epithelial subpopulations. Among immune-associated markers, transcriptomic expression of myeloid markers such as CD163 closely mirrored the increase in myeloid cell abundance detected by spatial analyses, whereas this trend was not observed at the proteomic level (Figures S2I and S2J). Conversely, proteomic expression of fibroblast markers such as COL1A2 was consistent with spatial and single-cell analyses, whereas corresponding transcriptomic changes were less apparent (Figures S2K and S2L). Despite these modality-specific differences, bulk transcriptomic and proteomic data broadly supported the cellular ecosystem remodeling revealed by spatial multi-omics analyses, highlighting dynamic changes in the composition and state of the breast tumor microenvironment during disease progression.

### 3 Basal cell depletion and epithelial barrier disruption in early breast cancer development

We next investigated dynamic changes in epithelial cell states during breast cancer progression. Re-clustering of epithelial populations from CosMx and CODEX datasets generated UMAP embeddings and expression landscapes of epithelial subpopulations (Figures 3A-3D). In the CosMx dataset, epithelial cells were classified into 10 functional subclusters, whereas 7 subclusters were identified in CODEX. Subcluster annotation was based on canonical marker genes and prior studies, defining luminal and basal lineages and their progenitor and mature states^27,28^. In the CosMx dataset, luminal progenitor cells were characterized by intermediate *GATA3* expression and absence of *PIP*, *XBP1* and *KRT5* expression, whereas mature luminal cells co-expressed *GATA3*, *PIP*, *XBP1* and *AZGP1*. Basal progenitor cells were defined by *KRT5* positivity and lack of *ACTA2* and *TAGLN*, while mature basal cells co-expressed *KRT5*, *ACTA2* and *TAGLN*. In the CODEX dataset, luminal progenitor cells were Keratin8/18-positive, Keratin5/14-negative, and CD66-positive (typically ER-negative), whereas mature luminal cells were Keratin8/18-positive, Keratin5/14-negative, and CD66-negative (typically ER-positive). Basal cells were defined by Keratin5/14 positivity, Keratin8/18 negativity, and expression of SMA and Caveolin.

**Figure 3.**
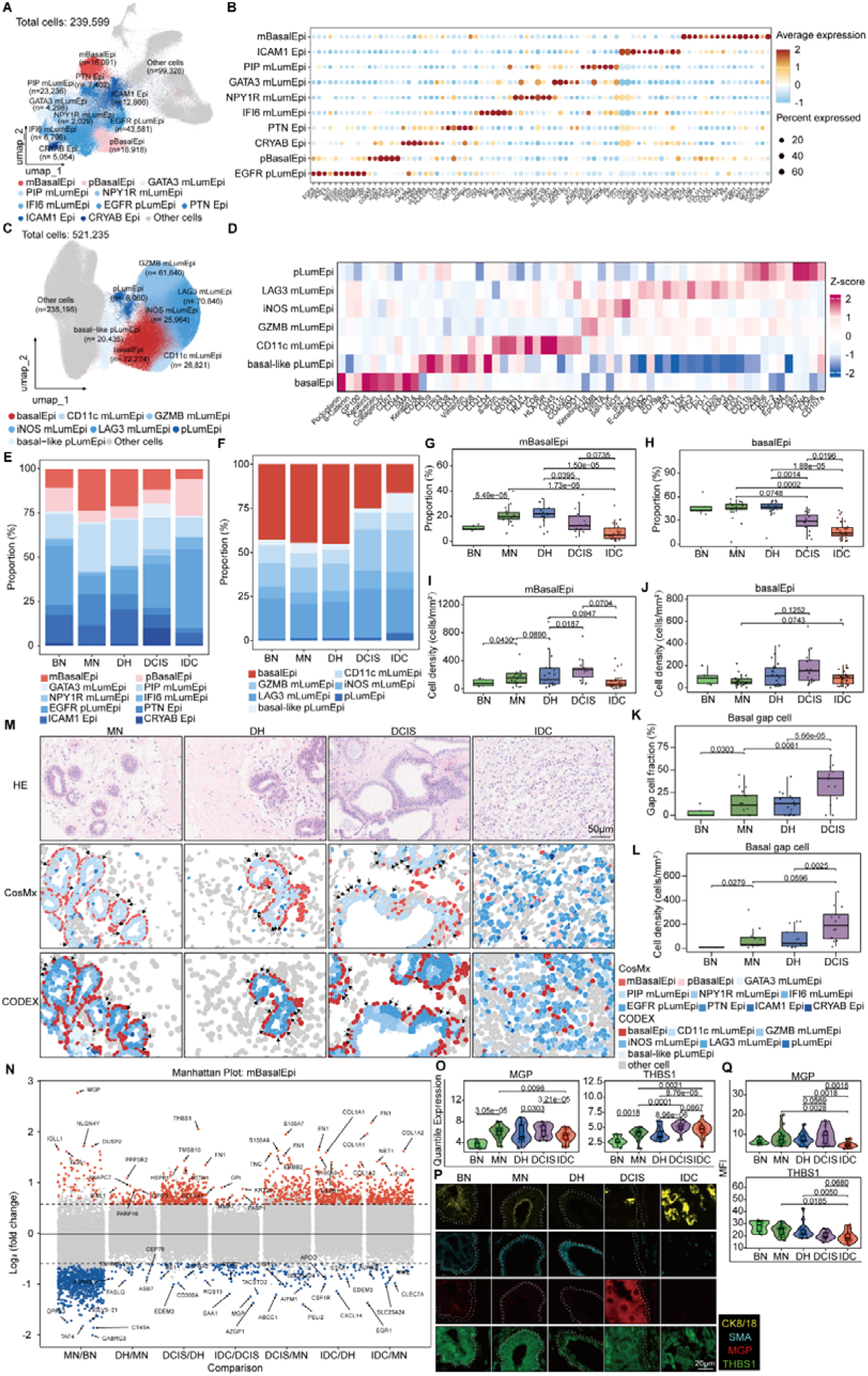
Early basal cell loss and barrier disruption during breast cancer progression. (A) UMAP projection of 239,599 cells from CosMx data, colored by epithelial subclusters; non-epithelial cells are shown in gray. (B) Dot plot showing average expression level (color) and percentage of expressing cells (size) of marker genes for each epithelial subcluster in CosMx data. (C) UMAP projection of 521,235 cells from CODEX data, colored by epithelial subclusters; non-epithelial cells are shown in gray. (D) Heatmap of Z-score normalized marker protein expression across epithelial subclusters in CODEX data. (E) Stacked bar plot showing the proportion of epithelial subclusters across BN, MN, DH, DCIS and IDC stages in CosMx data. (F) Stacked bar plot showing the proportion of epithelial subclusters across BN, MN, DH, DCIS and IDC stages in CODEX data. (G) Box-and-scatter plots depicting the proportion of mBasalEpi subcluster across disease stages in CosMx data. (H) Box-and-scatter plots depicting the proportion of basalEpi subcluster across disease stages in CODEX data. (I) Box-and-scatter plots showing cell density (cells/mm^2^) of mBasalEpi subcluster across disease stages in CosMx data. (J) Box-and-scatter plots showing cell density (cells/mm^2^) of basalEpi subcluster across disease stages in CODEX data. (K) Box-and-scatter plots depicting the proportion of basal gap cells in total basal cells across disease stages in CosMx data. (L) Box-and-scatter plots showing cell density (cells/mm^2^) of basal gap cells across disease stages in CosMx data. (M) Representative H&E, CosMx and CODEX maps of basal epithelial distribution; non-epithelial cells are shown in gray, black arrows indicate basal gap cells. Scale bars, 50 μm. (N) Manhattan plot of differentially expressed genes (DEGs) in mBasalEpi subcluster across pairwise disease stage comparisons; significance threshold: |log_2_(fold change)| >log_2_(1.5) and *P* < 0.05. (O) Violin-box-and-scatter plots showing the expression distribution of MGP and THBS1 in mBasalEpi subcluster across disease stages. (P) Representative immunofluorescence images of mIHC across MN, DH, DCIS and IDC stages, showing the expression of KRT18/8, SMA, MGP and THBS1. Scale bar, 20 μm. Two white dashed lines delineate the basal barrier. (Q) Violin-box-and-scatter plots showing the expression distribution of MGP and THBS1 in SMA+ cells across disease stages. Statistical analysis: Stage comparisons and statistical analyses were performed as described in Figure 2. All tests were performed using the t test for G-L, DEseq2 for N and O, limma for Q. *P* values are presented to four significant digits; *P <* 0.0001 is indicated in scientific notation. BN, normal from benign patient; MN, normal from malignant patient; DH, intraductal hyperplasia; DCIS, ductal carcinoma in situ; IDC, invasive ductal carcinoma; MFI, mean fluorescence intensity.

We observed profound remodeling of epithelial lineages during tumor progression. The proportion of mBasalEpi cells in CosMx and BasalEpi cells in CODEX progressively decreased from MN to IDC, with the most pronounced reduction during the transition from DH to DCIS (Figures 3E-3H). Previous studies have reported loss of myoepithelial cells and basement membrane integrity in DCIS^29^. Our spatial data further indicate that this process is already initiated during the DH-to-DCIS transition. In contrast, EGFR-positive luminal progenitor-like cells in CosMx and pLumEpi populations in CODEX (including basal-like pLumEpi and pLumEpi subtypes) increased progressively across disease stages, whereas mature luminal populations remained relatively stable (Figures 3E, 3F, and S3A-S3D). These findings are consistent with bulk transcriptomic data showing increasing KRT8 and decreasing KRT5 expression, supporting heterogeneous and asynchronous remodeling within epithelial compartments. Although basal epithelial cell proportions decreased, their absolute cell density increased progressively from MN to DCIS (Figures 3I and 3J). This divergence between proportion and density suggests that basal cell reduction is not absolute but instead reflects relative dilution driven by luminal expansion and ductal enlargement.

To quantify focal basal discontinuities, two investigators independently annotated basal-gap regions based on focal interruptions in basal-cell coverage and ductal morphology. Disagreements were resolved by consensus, and epithelial cells located within the concordantly annotated regions were designated basal gap cells. Both the proportion and spatial density of basal gap cells increased during disease progression, with the most prominent changes observed during the DH-to-DCIS transition (Figures 3K, 3L, S3E, and S3F). Quantification of ductal structures across fields of view revealed no significant increase in ductal number across stages (Figure S3G), indicating that the expansion of basal gap cells is more likely associated with ductal dilation and loss of basal continuity rather than increased glandular abundance. Consistently, H&E staining and spatial imaging from both platforms revealed progressive ductal expansion, luminal protrusion, and loss of basal epithelial continuity from MN through DH, DCIS, and IDC (Figure 3M). Analysis of CODEX raw images further demonstrated that KRT14 continuity was already disrupted in DH lesions, while Vimentin and SMA showed more pronounced focal loss in DCIS, indicating a progressive deterioration of basal structural integrity that intensifies during the DH-to-DCIS transition (Figure S3H).

To investigate potential molecular mechanisms underlying basal layer disruption, we performed differential expression analysis of mBasalEpi cells across disease stages (Figures 3N, 3O, S3I-S3K, Table S3). Compared with BN, MN samples showed upregulation of *MGP* and *NLGN4Y* and downregulation of *GABRG3*. In the MN-to-DH transition, *PPP3R2*, *ANAPC7* and *IFNL1* were upregulated, whereas *FASLG*, *AURKB* and *ASB7* were downregulated. Notably, the largest number of differentially expressed genes was observed between DH and DCIS. During this transition, basal cells exhibited strong extracellular matrix and invasion-associated transcriptional reprogramming, including increased expression of *THBS1*, *TMSB10* and *FN1*. Selected genes with the most pronounced changes were further validated using mIHC. Cells in the top 10% of SMA or KRT8/18 expression were defined as positive populations. MGP protein levels increased markedly from DH to DCIS, whereas GABRG3 decreased, consistent with spatial transcriptomic results. In contrast, THBS1 protein levels showed an opposite trend, decreasing during DH-to-DCIS progression despite increased transcript abundance (Figures 3P, 3Q, S3L, and S3M). This discordance between transcript and protein levels is unlikely to reflect technical artifacts, as extracellular matrix (ECM)-related secreted proteins are known to exhibit weak or even negative mRNA-protein correlations^30^.

Finally, we validated these observations using bulk multi-omics datasets. Basal membrane-associated proteins were significantly reduced in IDC (Figure S3N). Consistent with spatial analyses, THBS1 showed a progressive increase in both transcriptomic and proteomic datasets from MN to DCIS, whereas MGP protein levels also increased during this transition (Figures S3O and S3P). Collectively, these results demonstrate progressive disruption of the basal cell layer during the transition from DH to DCIS, accompanied by marked transcriptional remodeling of basal epithelial cells, suggesting that these features may serve as potential biomarkers for predicting malignant transformation risk in DH lesions.

### 4 Multi-omics characterization of invasion-associated features in early breast cancer development

We first performed gene set enrichment analysis (GSEA) of mutation-associated genes across disease stages. Enrichment of ECM-receptor interaction was observed during the MN-to-DH transition (Figure S4A). The DH-to-DCIS transition showed broader activation of ECM-receptor interaction, focal adhesion, integrin signaling, and other ECM-related pathways (Figure 4A). These findings indicate progressive remodeling of ECM- and adhesion-associated networks during the transition from DH to DCIS. Analysis of genes within the ECM-receptor interaction pathway revealed frequent somatic mutations in components of the PI3K–AKT signaling axis. Notably, PIK3CA mutations were more frequent in IDC than in DCIS, suggesting progressive accumulation of PI3K–AKT pathway alterations during tumor evolution. Importantly, PIK3CA mutations were already detectable in a subset of DH and DCIS samples, indicating that activation of this pathway may occur early in tumorigenesis (Figure 4B). Consistently, GSEA of the DH-to-DCIS transition also demonstrated significant enrichment of ECM-related programs (Figure 4C).

**Figure 4.**
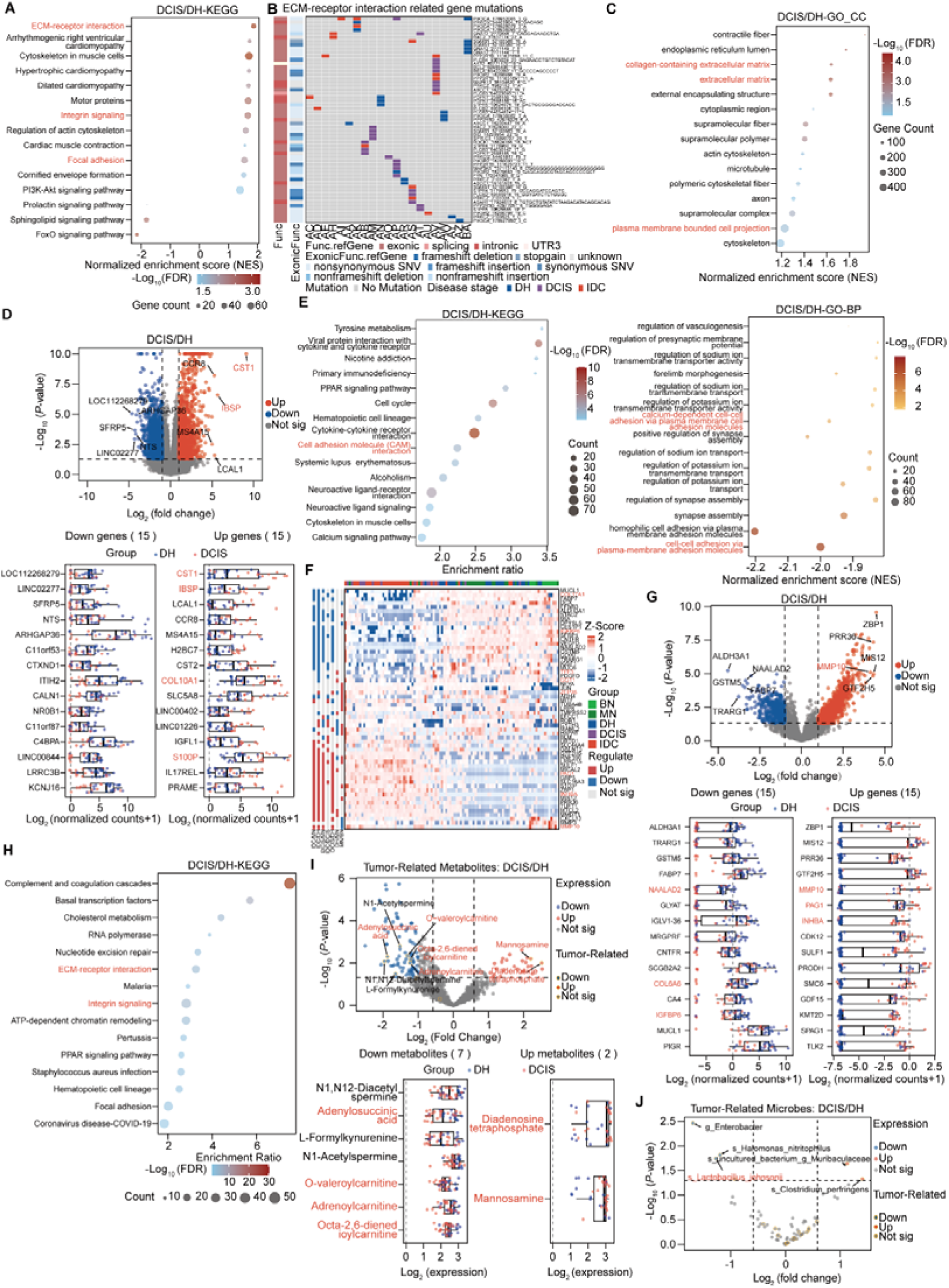
Multi-omics characterization of invasion-associated features in early breast cancer development. (A) Bubble plot showing enriched KEGG pathways from GSEA for somatic mutations in DCIS compared to DH. Dot size indicates gene count, and color represents -log_10_(FDR) (blue to red, significant) (B) Mutation heatmap of ECM-receptor interaction pathway-enriched genes during disease progression. Based on the enriched genes of the ECM-receptor interaction pathway identified in (A), this heatmap shows the mutation status of ECM-related genes in patients at different disease stages. (C) Bubble plot showing the enrichment results of GO cellular component (CC) terms by GSEA for somatic mutations in the DH vs MN comparison. (D) Box-and-volcano plots showing bulk transcriptomic changes during the DH-to-DCIS transition. The volcano plot (upper) compares bulk transcriptomic expression between DCIS and DH, with the top 5 up- and down-regulated genes labeled. Boxplots (lower) display expression distributions [log_2_(normalized count + 1)] of the top 15 up- and down-regulated genes ranked by |log_2_FC|. (E) Pathway enrichment plots showing bulk transcriptomic changes during the DH-to-DCIS transition. Both the KEGG enrichment analysis bubble plot (left) and the GSEA dot plot for GO biological process (BP) terms (right) revealed significant enrichment of invasion-related pathways. (F) Heatmap showing z-score normalized expression levels of the top 5 upregulated and top 5 downregulated proteins (|FC| > 2, *P* < 0.05) across BN, MN, DH, DCIS, and IDC stages. (G) Box-and-volcano plots showing bulk proteomic changes during the DH-to-DCIS transition. The volcano plot (upper) compares bulk protein abundance between DCIS and DH, with the top 5 up- and down-regulated proteins labeled. Boxplot (lower) displays expression distribution [log_2_ (normalized count + 1)] of the top 15 up- and down-regulated proteins ranked by |log_2_FC|. (H) Bubble plot showing KEGG pathway enrichment analysis of differentially expressed proteins between DCIS and DH stages. (I) Box-and-volcano plots showing bulk metabolomic changes during the DH-to-DCIS transition. Volcano plots (upper) compare metabolite abundance between DCIS and DH stages, with invasion-associated metabolites features highlighted by yellow borders and labeled. Box plots (lower) show the expression patterns of invasion-associated metabolites. (J) Volcano plots showing bulk microbiome changes during the DH-to-DCIS transition. Statistical analysis: Stage comparisons and statistical analyses were performed as described in Figure 2. For differential expression analysis in volcano plots of D, G, I and J, significance was defined as P value < 0.05 and |log_2_(fold change)| > 1. All enrichment analyses were performed using the clusterProfiler package. KEGG enrichment analysis was performed using the enrichKEGG function with Benjamini-Hochberg correction, applying a significance threshold of FDR < 0.1 for E left and H. GSEA analysis was performed using the gseKEGG, gseGO (CC), and gseGO (BP) functions, with a significance threshold of FDR < 0.05 for A, C, and E right. DH, intraductal hyperplasia; DCIS, ductal carcinoma in situ.

Unsupervised QC-UMAP analysis of bulk transcriptomes showed clear separation of samples across disease stages. Malignant samples (DCIS and IDC) were distinct from MN samples, whereas DH samples occupied an intermediate position along a continuous trajectory, suggesting acquisition of early malignant-like molecular features (Figure S4B). Differential expression analysis identified 286, 2,557, and 889 differentially expressed genes across MN-to-DH, DH-to-DCIS, and DCIS-to-IDC transitions, respectively, with the largest transcriptional shift occurring during DH-to-DCIS progression (Figure S4C). In this transition, invasion-associated genes such as CST1, S100P, COL10A1, and IBSP were significantly upregulated (Figure 4D). Pathway analysis further highlighted altered cell adhesion molecule (CAM) interactions and enrichment of cell–cell adhesion programs during DH-to-DCIS progression (Figure 4E).

Proteomic profiling similarly distinguished malignant from non-malignant stages (Figure S4D), with DCIS and IDC samples showing increased numbers of detectable proteins compared with other stages (Figure S4E). Differential protein analysis identified the most pronounced changes during the DH-to-DCIS transition (1,579 proteins), predominantly characterized by upregulation (Figure S4F). Several invasion-associated proteins exhibited dynamic stage-specific patterns (Figure 4F). Compared with MN, DH lesions showed increased expression of GGT1, TMPRSS2, and TFF3, which have been implicated in tumor invasion, migration, or abnormal epithelial differentiation, whereas ZFP36, a known suppressor of inflammatory and oncogenic transcriptional programs, was downregulated (Figure S4G). At the DCIS stage, proteins involved in extracellular matrix degradation, EMT, and migration regulation, including MMP10, PAG1, and INHBA, were further upregulated compared with DH. In contrast, COL6A6, IGFBP6, and NAALAD2, associated with reduced invasive potential or loss of differentiation, were downregulated (Figure 4G). In IDC, COL17A1 was decreased, suggesting weakened basement membrane attachment and enhanced invasive capacity (Figure S4H). Pathway analysis confirmed significant enrichment of ECM-receptor interaction, integrin signaling, and focal adhesion pathways, together with activation of basement membrane organization programs (Figures 4H and S4I, S4J).

Metabolomic profiling revealed the largest number of differential metabolites during the DH-to-DCIS transition (n = 77) (Figure S4K). Increased levels of mannose-related metabolites and diadenosine tetraphosphate suggested enhanced glycometabolism and nucleotide metabolism, whereas multiple carnitines and energy metabolism-related metabolites were decreased, indicating a metabolic shift from homeostasis toward reprogramming (Figure 4I). Exploratory 16S rRNA gene profiling identified a lower relative abundance of ASVs assigned to Lactobacillus johnsonii in DCIS than in DH, suggesting depletion of potentially protective microbial populations and disruption of local tumor-suppressive microecological balance (Figure 4J).

Collectively, integrative multi-omics analyses across genomic, transcriptomic, proteomic, metabolomic, and microbiome consistently identified the DH-to-DCIS transition as a critical window characterized by extensive activation of invasion-associated programs. These results support a model in which matrix remodeling and cell adhesion reprogramming are already initiated prior to histologically defined DCIS.

### 5 Microenvironmental regulation of basal cell behavior during the initiation of breast cancer invasion

Given the marked spatial redistribution and transcriptional alterations of mBasalEpi cells during the DH-to-DCIS transition, we next investigated intercellular communication networks linking epithelial subpopulations across disease progression. Ligand-receptor interaction analysis of CosMx and CODEX datasets revealed a marked increase in signaling between EGFR-positive luminal progenitor-like cells (EGFR⁺ pLumEpi) and mBasalEpi cells during DH and DCIS, reaching maximal intensity at the DCIS stage (Figure 5A; Table S4A). This suggests that EGFR⁺ luminal-to-basal communication is initiated in DH lesions and further amplified during DCIS progression. To define the molecular basis of this enhanced interaction, we examined the top 20 ligand–receptor pairs mediating EGFR⁺ pLumEpi–mBasalEpi crosstalk across disease stages (Figures 5B, 5C; Figure S5A). Among these, SEMA4D-ERBB2 interactions predominantly reflected signaling from EGFR⁺ pLumEpi toward mBasalEpi cells. SEMA4D, a classical axon guidance and migration-associated molecule, has been implicated in tumor invasion and microenvironmental remodeling^31,32^, whereas ERBB family signaling plays a central role in mammary epithelial proliferation and malignant transformation. Mechanistically, SEMA4D can activate ERBB2 signaling through Plexin-mediated pathways^32–34^. THBS1-SDC1 interactions were primarily directed from mBasalEpi cells to EGFR⁺ pLumEpi cells (Figure 5C; Figure S5A). THBS1, a key extracellular matrix and adhesion-related molecule^35^, was highly enriched in mBasalEpi cells, consistent with our prior observations. This indicates that basal cells are not merely passive structural components but may actively transmit ECM- and adhesion-related signals to regulate adjacent luminal-like populations^29^.

**Figure 5.**
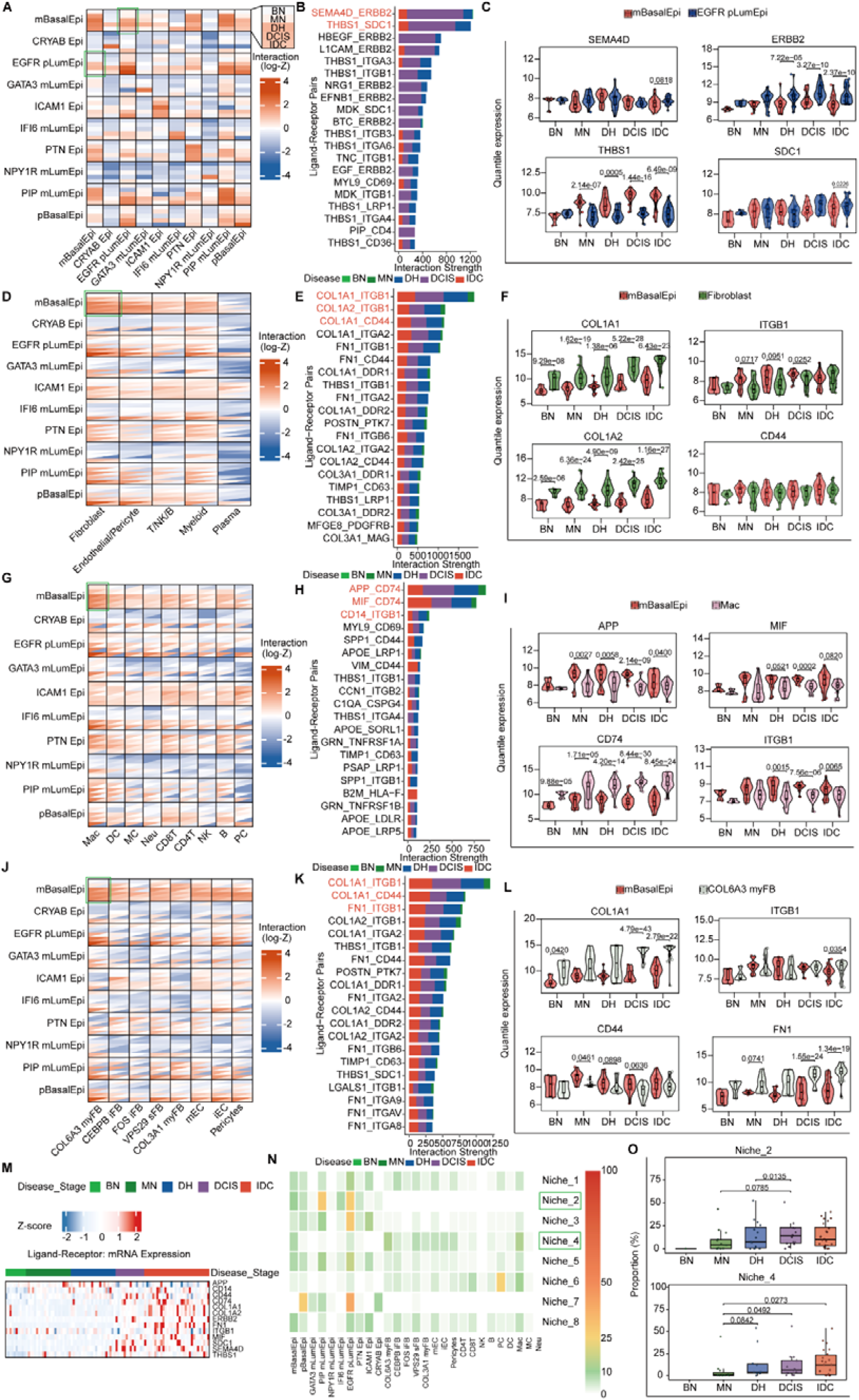
Microenvironmental regulation of basal cell behavior during the initiation of breast cancer invasion. (A) Heatmap showing ligand–receptor interaction strength between epithelial subclusters in CosMx data. Columns represent sender cells and rows represent receiver cells. The green box highlights interactions from EGFR pLumEpi to mBasalEpi. Each interaction is stratified by disease stage (top to bottom: BN, MN, DH, DCIS, and IDC), with color indicating normalized counts of significant interactions. (B) Horizontal bar plot showing the top 20 ligand-receptor interaction pairs between mBasalEpi and EGFR pLumEpi, ranked by interaction strength. (C) Violin-box-and-scatter plots depicting the expression levels of SEMA4D, ERBB2, THBS1 and SDC1 in mBasalEpi and EGFR pLumEpi across disease stages. (D) Heatmap showing ligand-receptor interaction strength between epithelial subclusters and major cell types in CosMx data. The green box highlights interactions between mBasalEpi and fibroblasts. For each cell pair, interactions above the diagonal represent signaling from row cell populations to column cell populations, whereas interactions below the diagonal represent signaling from column cell populations to row cell populations. Color indicates normalized counts of significant interactions. (E) Horizontal bar plot showing the top 20 ligand-receptor interaction pairs between mBasalEpi and fibroblasts, ranked by interaction strength. (F) Violin-box-and-scatter plots depicting the expression levels of COL1A1, ITGB1, COL1A2 and CD44 in mBasalEpi and fibroblasts across disease stages. (G) Heatmap of ligand-receptor interaction strength between epithelial subclusters and myeloid subclusters in CosMx data; the green box highlights interactions between mBasalEpi and macrophages (Mac). Upper and lower triangles indicate opposite signaling directions, as described in Figure 5D. (H) Horizontal bar plot showing the top 20 ligand-receptor interaction pairs between mBasalEpi and macrophages, ranked by interaction strength. (I) Violin-box-and-scatter plots depicting the expression levels of APP, MIF, CD74 and ITGB1 in mBasalEpi and macrophages across disease stages. (J) Heatmap of ligand-receptor interaction strength between epithelial subclusters and myofibroblast subclusters in CosMx data; the green box highlights interactions between mBasalEpi and COL6A3 myFB. Upper and lower triangles indicate opposite signaling directions, as described in Figure 5D. (K) Horizontal bar plot showing the top 20 ligand-receptor interaction pairs between mBasalEpi and COL6A3 myFB, ranked by interaction strength. (L) Violin-box-and-scatter plots depicting the expression levels of COL1A1, ITGB1, CD44 and FN1 in mBasalEpi and COL6A3 myFB across disease stages. (M) Heatmap of basal-centered cell-cell interaction genes expression in bulk transcriptome. (N) Heatmap of spatial neighborhood cell composition in CosMx data; rows represent niche types, columns represent cell subclusters, and yellow boxes highlight Niche_2 and Niche_4. (O) Box-and-scatter plots showing the proportion of Niche_2 and Niche_4 across disease stages. Statistical analysis: Stage comparisons and statistical analyses were performed as described in Figure 2. All tests were performed using the limma for C, F, I, L, and t test for O. *P* values are presented to four significant digits; *P <* 0.0001 is indicated in scientific notation. BN, normal from benign patient; MN, normal from malignant patient; DH, intraductal hyperplasia; DCIS, ductal carcinoma in situ; IDC, invasive ductal carcinoma.

We next extended the analysis to epithelial-stromal communication. Interaction strength between mBasalEpi cells and both fibroblasts and myeloid cells progressively increased from MN to DH and remained elevated in DCIS (Figure 5D; Table S4B), suggesting early activation of stromal–basal crosstalk during tumor initiation. Fibroblast–mBasalEpi interactions were dominated by ECM-related ligand–receptor pairs, including COL1A1-ITGB1, COL1A2-ITGB1, and COL1A1-CD44 (Figures 5E, 5F; Figure S5B). Type I collagen (COL1A1/COL1A2), a major fibroblast-derived ECM component^36^, signals through ITGB1, a key receptor for mechanical sensing and adhesion maintenance in basal epithelial cells^37^. These interactions suggest that fibroblast-derived collagen matrices may alter basal cell adhesion and cytoskeletal organization through integrin signaling, thereby compromising epithelial barrier stability. In addition, COL1A1–CD44 interactions, which regulate cell migration and mechanotransduction^38^, were also enhanced, suggesting additional CD44-mediated modulation of basal cell state.

To further dissect immune involvement, we reclassified immune populations from CosMx and CODEX datasets into 9 and 11 subclusters, respectively (Figures S5C–S5F). Macrophage-mBasalEpi interactions were significantly increased as early as the DH stage and persisted throughout disease progression (Figure 5G; Table S4C), indicating early activation of macrophage-basal communication prior to overt invasion. Macrophages expanded progressively during tumor evolution, with both abundance and spatial density increasing from MN to IDC (Figures S5G-S5L), consistent with the accumulation of tumor-associated macrophages reported in breast cancer progression^39,40^. These findings suggest that macrophage–basal communication is an early event rather than a consequence of invasive transformation. Ligand–receptor analysis revealed bidirectional signaling between macrophages and mBasalEpi cells. APP-CD74 and MIF-CD74 interactions were predominantly directed from mBasalEpi cells to macrophages, suggesting that basal cells actively participate in shaping inflammatory and stress-related immune programs. In contrast, CD14-ITGB1 interactions were mainly macrophage-to-basal, indicating that macrophages may regulate basal cell state through integrin-associated signaling (Figure 5H; Figure S5M).

We next analyzed stromal heterogeneity by re-clustering mesenchymal populations from CosMx and CODEX datasets, identifying 8 and 10 subclusters, respectively (Figures S6A-S6D). Interaction analysis showed that COL6A3_myFB and COL3A1_myFB exhibited the strongest communication with mBasalEpi cells during DH and DCIS (Figure 5J; Table S4D), implicating myofibroblast activation in basal layer remodeling. Consistently, myofibroblast-associated populations progressively expanded during disease progression, with both proportion and spatial density increasing across stages (Figures S6E-S6J), highlighting stromal activation as a hallmark of tumor evolution. Further ligand-receptor analysis of the COL6A3_myFB-mBasalEpi revealed dominant ECM-mediated interactions, including COL1A1-ITGB1, COL1A1-CD44, and FN1-ITGB1 (Figures 5K, 5L, Figure S6K). These signals converge on integrin-mediated adhesion and mechanotransduction pathways, suggesting that activated stromal cells remodel basal cell adhesion and tissue architecture through ECM deposition. Bulk transcriptomic analysis further confirmed that genes involved in these ligand-receptor interactions were significantly enriched in DCIS and IDC samples (Figure 5M).

Finally, spatial niche analysis identified eight major microenvironmental niches. mBasalEpi cells were predominantly enriched in niche_2 and niche_4. Niche_2 was characterized by EGFR⁺ pLumEpi and PIP⁺ mature luminal cells, whereas niche_4 was enriched for COL6A3_myFB and macrophages (Figure 5N). Consistently, both niche_2 and niche_4 increased during tumor progression, with significant expansion during the DH-to-DCIS transition (Figure 5O; Figure S5L). These results suggest that EGFR⁺ pLumEpi cells, myofibroblasts, and macrophages constitute coordinated microenvironmental hubs that converge on basal cell remodeling during early breast cancer invasion.

### 6 Luminal leader cells breach the basal cell layer to generate gaps

We quantified the cellular composition of Basal gap regions across disease stages to identify epithelial populations contributing to basal discontinuity. EGFR⁺ pLumEpi cells showed the most pronounced enrichment within Basal gap regions during disease progression in CosMx (Figures 6A, 6B). Consistent results were obtained in the CODEX dataset, where pLumEpi cells represented the dominant luminal population expanding within Basal gap regions (Figures 6C, 6D). These findings suggest that basal layer disruption is not solely driven by alterations in myoepithelial cells^15^ but is strongly associated with a specific EGFR⁺ luminal subpopulation, identifying a candidate epithelial state associated with basal barrier discontinuity.

**Figure 6.**
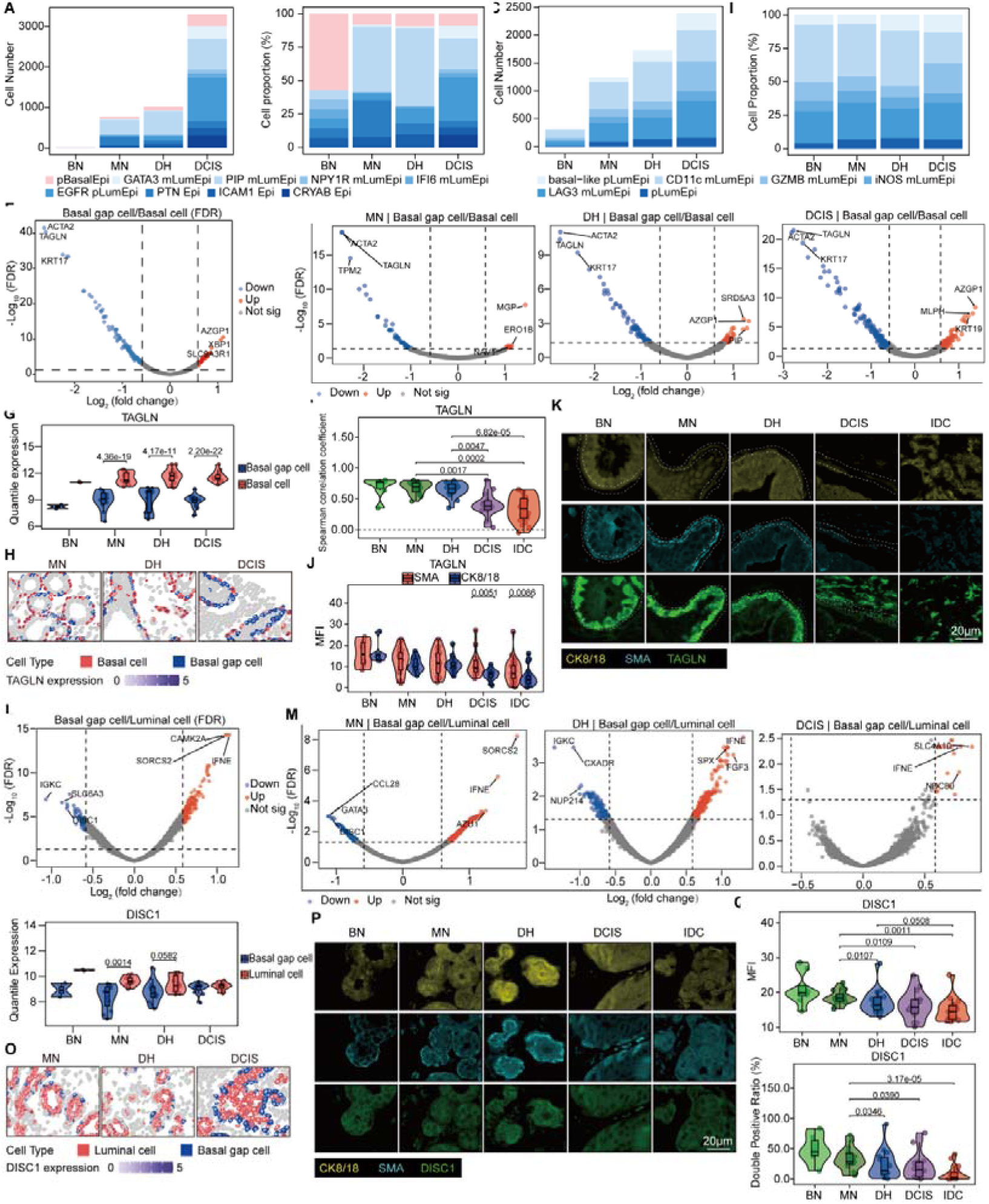
Luminal leader cells breach the basal cell layer to generate gaps. (A) Bar plot showing the cell number of epithelial subclusters at basal gap regions in CosMx data. (B) Stacked bar plot showing the cell proportion of epithelial subclusters at basal gap regions in CosMx data. (C) Bar plot showing the cell number of epithelial subclusters at basal gap regions in CODEX data. (D) Stacked bar plot showing the cell proportion of epithelial subclusters at basal gap regions in CODEX data. (E) Volcano plot of DEGs between basal gap cells and basal cells across all disease stages in CosMx data. (F) Volcano plots of DEGs between basal gap cells and basal cells stratified by disease stage (MN, DH, DCIS) in CosMx data. (G) Violin-box-and-scatter plots depicting the expression levels of representative DEGs (TAGLN) in basal gap cells and basal cells across disease stages; *P* values represent FDR from differential analysis. (H) Representative spatial expression maps of the above DEGs in basal gap cells and basal cells across disease stages, with cell type indicated by border color and gene expression level represented by fill intensity. (I) Violin-box-and-scatter plots depicting the spearman correlation coefficient of protein TAGLN in SMA+ cells across disease stages. (J) Violin-box-and-scatter plots depicting the MFI of protein TAGLN in SMA+ cells and CK8/18+ cells across disease stages. (K) Representative immunofluorescence images of mIHC across disease stages, showing the expression of CK8/18, SMA and TAGLN. Scale bar, 20 μm. Two white dashed lines delineate the basal barrier. (L) Volcano plot of DEGs between basal gap cells and all luminal cells across all disease stages in CosMx data. (M) Volcano plots of DEGs between basal gap cells and luminal cells stratified by disease stage (MN, DH, DCIS) in CosMx data. (N) Violin-box-and-scatter plots depicting the expression levels of representative DEGs (DISC1) in basal gap cells and luminal cells across disease stages; *P* values represent FDR from differential analysis. (O) Representative spatial expression maps of the above DEGs in basal gap cells and luminal cells across disease stages, with cell type indicated by border color and gene expression level represented by fill intensity. (P) Representative immunofluorescence images of mIHC across disease stages, showing the expression of CK8/18, SMA and DISC1. Scale bar, 20 μm. (Q) Violin-box-and-scatter plots depicting the MFI of protein DISC1 in CK8/18^+^ cells across disease stages. (R) Violin-box-and-scatter plots depicting the double positive ratio of protein DISC1 in CK8/18^+^ cells across disease stages. Statistical analysis: For DEG identification in volcano plots, the significance cutoff was set as FDR < 0.05 and |log_2_(fold change)| > log_2_(1.5). Stage comparisons and statistical analyses were performed as described in Figure 2. All tests were performed using limma for E, F, G, J, L, M and N, and t tests for I, Q and R. *P* values are presented to four significant digits; *P* < 0.0001 is indicated in scientific notation. BN, normal from benign patient; MN, normal from malignant patient; DH, intraductal hyperplasia; DCIS, ductal carcinoma in situ; IDC, invasive ductal carcinoma; MFI, mean fluorescence intensity.

To define the molecular features of Basal gap cells, we next compared basal gap cells with basal epithelial cells. CODEX revealed marked downregulation of basal markers, including SMA, KRT5 and KRT14, in basal gap regions (Figures S7A-S7C). CosMx analysis further showed consistent suppression of myoepithelial-associated genes, including *KRT17*, *ACTA2* and *TAGLN*, whereas genes involved in secretion, stress response, and membrane signaling, including *AZGP1*, *XBP1*, and SLC9A3R1, were significantly upregulated (Figure 6E). These alterations were most pronounced during the DH and DCIS stages (Figure 6F). Quantitative analysis confirmed increased *AZGP1* and *MLPH* expression and decreased *TAGLN* expression in basal gap cells relative to basal epithelial cells (Figure 6G; Figure S7D). Spatial mapping further demonstrated enrichment of *AZGP1* and *MLPH* within basal gap regions, whereas *TAGLN* was restricted to intact basal layers (Figure 6H; Figure S7E). Multiplex immunohistochemistry validation confirmed a progressive reduction in the correlation between TAGLN, MLPH, AZGP1 and SMA-positive (SMA^+^) cells during disease progression. TAGLN also showed higher fluorescence intensity within SMA^+^ cells (Figures 6I-6K; Figure S7F).

Given that basal gap regions are predominantly filled by luminal cells, we next compared basal gap cells with other luminal populations to identify features associated with gap formation. Basal gap cells exhibited reduced expression of KRT8/18 from the MN stage, with further decreases in ER, KRT8/18, and pan-cytokeratin observed during the DH stage, indicating early alterations in epithelial differentiation states in CODEX (Figures S7G-S7I). Spatial transcriptomic analysis further revealed that basal gap cells were characterized by upregulation of *IFNE*, *FGF3* and *SLC4A10*, and downregulation of *IGKC*, *DISC1* and *SLC6A3* compared with other luminal cells (Figures 6L, 6M). IFNE, FGF3 and SLC4A10 are associated with secretory activity and microenvironmental regulation^41,42^, suggesting that Basal gap cells represent a distinct luminal state with active paracrine and niche-modulating functions that may contribute to local basal disruption. The most extensive molecular reprogramming of basal gap cells occurred during the DH stage, whereas both the number of differentially expressed genes and proteins decreased in DCIS. This suggests that invasion-associated molecular events are initiated during DH and that the structural abnormalities observed in DCIS likely reflect the cumulative outcome of earlier molecular remodeling. DISC1 was highly expressed in luminal cell populations rather than basal gap cells (Figures 6N, 6O). Consistently, DISC1 was predominantly expressed in KRT8/18-positive (KRT8/18+) cells (Figure 6P). Moreover, both the mean fluorescence intensity and the positive proportion of DISC1 in KRT8/18⁺ cells progressively decreased with disease progression (Figures 6Q, 6R). Finally, bulk transcriptomic analysis showed that DISC1 expression remained relatively stable during the MN-to-DH transition but was significantly reduced in DCIS (Figure S7J), further supporting stage-specific regulation during early tumor progression.

### 7 Assessment of malignant transformation risk in benign breast lesions

To evaluate whether molecular alterations identified in DH-associated early invasion are associated with malignant transformation risk in benign breast lesions, we established an independent validation cohort. This cohort included 90 patients initially diagnosed with benign breast tumors who underwent surgical resection. All patients subsequently experienced an ipsilateral breast lesion at least six months later, classified as benign or malignant. Based on clinical outcomes, patients were classified into a benign-to-benign recurrence group and a benign-to-malignant transformation group. A total of 180 formalin-fixed paraffin-embedded (FFPE) samples from initial and recurrent surgeries were retrieved and assembled into tissue microarrays for mIHC analysis, together with comprehensive clinical annotation (Figure 7A). Marker expression within SMA+ cells differed significantly between the two groups at the initial benign stage. Compared with the first benign lesion (BF) group, first benign lesion from malignant patient (MF) group showed significantly decreased GABRG3 expression and increased expression of TAGLN, MLPH and AZGP1, as reflected by mean fluorescence intensity (Figures 7B, 7C). These differences suggest that the above molecules already exhibit detectable stratified expression patterns at the histologically benign stage and may serve as potential biomarkers for predicting subsequent malignant transformation risk.

**Figure 7.**
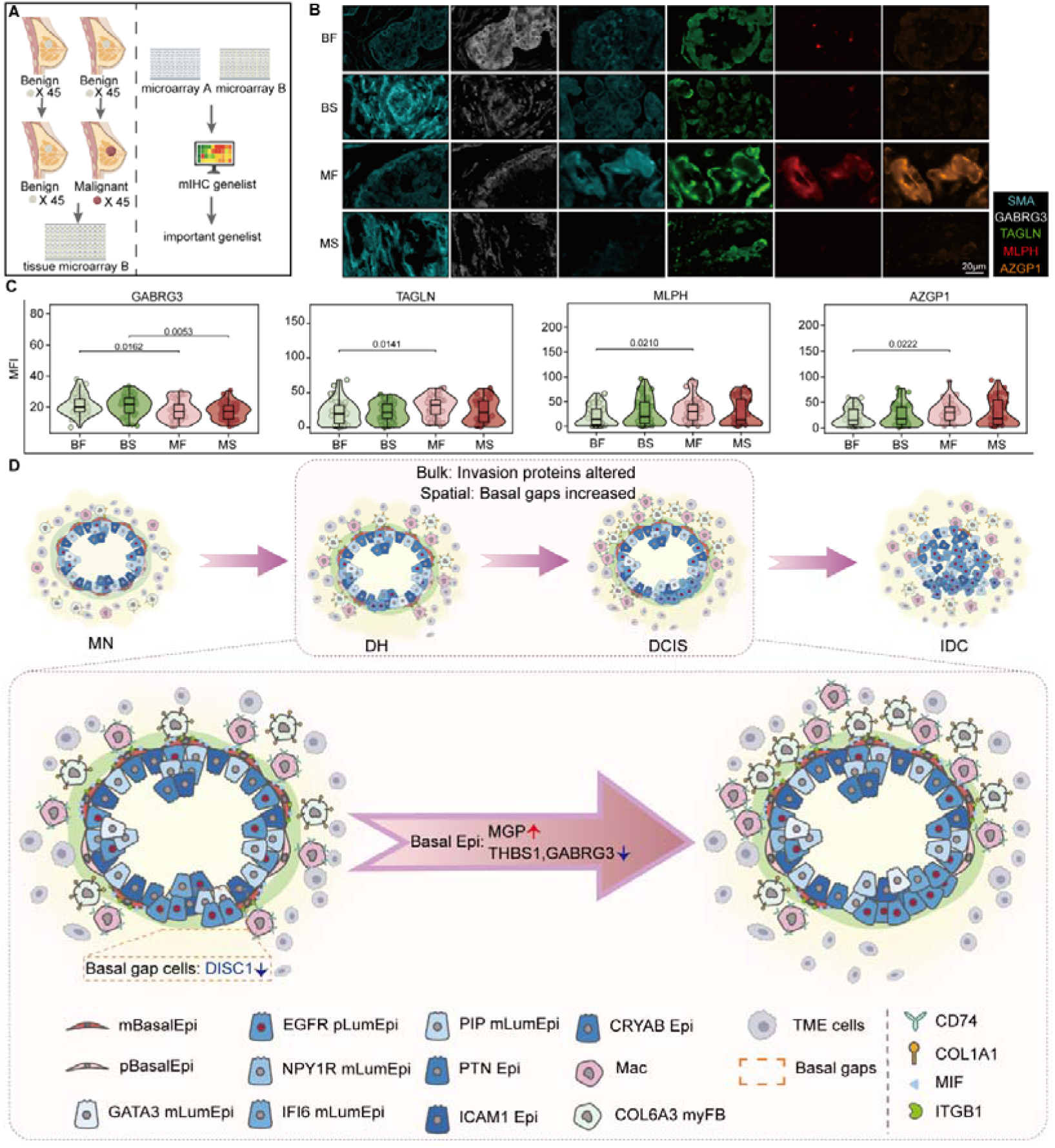
Assessment of malignant transformation risk in benign breast lesions. (A) Schematic illustrating the workflow of Cohort for risk assessment of malignant transformation in benign breast tumors. (B) Representative immunofluorescence images of mIHC across BF, BS, MF and MS samples, showing the expression of SMA, GABRG3, TAGLN, MLPH and AZGP1. Scale bar, 20 μm. (C) Violin-box-and-scatter plots depicting the mean fluorescence intensity of GABRG3, TAGLN, MLPH and AZGP1 in SMA+ cells across BF, BS, MF and MS groups. (D) Schematic model of basal barrier disruption during breast cancer progression. Statistical analysis: Stage comparisons and statistical analyses were performed as described in Figure 2. All tests were performed using the limma for C. *P* values are presented to four significant digits; *P* < 0.0001 is indicated in scientific notation. BF, first benign lesion; BS, second benign lesion; MF, first benign lesion from malignant patient; MS, second malignant lesion from malignant patient; MN, normal from malignant patient; DH, intraductal hyperplasia; DCIS, ductal carcinoma in situ; IDC, invasive ductal carcinoma.

## Discussion

This study profiles a human breast tissue cohort spanning the cross-sectional pathological stages MN, DH, DCIS and IDC. Using a space-for-time design, we compare spatial and molecular features across these stages within a clinically defined cohort. Unlike prior studies predominantly focused on invasive and metastatic disease^43,44^, we integrate single-cell spatial transcriptomics, spatial proteomics, and multi-layer bulk omics to map stage-associated patterns across cellular, tissue and system-level molecular networks, providing a framework for understanding early breast cancer initiation.

A central finding of this study is that oncogenic programs are already activated at the DH stage, prior to histologically defined DCIS and IDC. Although breast cancer has long been modeled as a linear progression from DH to DCIS to IDC, direct human evidence defining the timing of malignant initiation has been lacking. The basal/myoepithelial layer, a key structural and functional barrier maintaining ductal integrity, is widely recognized as a hallmark structure whose disruption enables invasion^45,46^. While focal basal layer loss has been reported in DCIS and associated with increased invasive risk^29,47^, our spatial multi-omics data demonstrate that loss of basal layer continuity predominantly emerges during the DH-to-DCIS transition. This process is accompanied by coordinated extracellular matrix remodeling, dysregulated adhesion signaling, and broad activation of invasion-associated programs. Importantly, the DH-to-DCIS transition exhibits more extensive transcriptomic, proteomic, and metabolomic reprogramming than the DCIS-to-IDC transition, suggesting that the major preparatory phase of malignant transformation occurs earlier than previously appreciated. This is consistent with evidence that most genomic evolution precedes DCIS^48^ and with reports that histologically normal breast tissue may already harbor cells with tumor-associated copy number alterations^49^. Together, these findings support a revised model in which invasion is not triggered at the moment of basement membrane breach, but is biologically primed during DH, with DCIS representing a progressive histological manifestation of an already established malignant trajectory.

Beyond temporal dynamics, our data reveal a cellular mechanism underlying basal barrier disruption. Basal cells have been recognized not only as structural components but also as key regulators of the epithelial microenvironment^50^. However, the identity of cells initiating basal layer penetration has remained unclear. Here, we identify an EGFR⁺ luminal progenitor-like population enriched at basal gap regions across the pathological stages examined. These cells localize specifically to sites of basal discontinuity, and their transcriptional reprogramming precedes detectable alterations in basal cells, peaking at the DH stage. Spatial interaction analyses further reveal a highly coordinated signaling network involving basal cells, EGFR⁺ luminal cells, macrophages, and COL6A3⁺ fibroblasts, indicating that basal barrier discontinuity is associated with luminal cell plasticity, basal cell state transition and microenvironmental remodeling (Figure 7D).

This mechanism aligns closely with emerging models of collective invasion^51,52^, where a subset of “leader cells” orchestrates tissue invasion^51–54^. We identify a candidate leader-like population enriched at regions of basal discontinuity in human tissues. Notably, transcriptional differences between candidate leader-like cells and other luminal cells are maximal at the DH stage but diminish markedly in DCIS, This transient nature is consistent with prior reports describing high plasticity of candidate leader-like cells states^55^. Candidate leader-like cells exhibit enrichment of immune regulatory, proliferative, and neuro-associated signaling pathways, supporting emerging concepts that neural and immune systems jointly modulate tumor invasion and progression^56^.

From a translational standpoint, these findings provide a conceptual framework for risk stratification of DH lesions. A major clinical challenge is distinguishing indolent DH lesions from those that will progress to DCIS or IDC. Our results demonstrate that malignant reprogramming is already detectable at the DH stage, defining a previously unrecognized window for early risk assessment. We further identify spatial molecular features associated with malignant initiation, including GABRG3, TAGLN, MLPH, and AZGP1, which are validated in an independent cohort with clinical outcome data. While further validation in large multicenter studies is required, our findings establish the feasibility of using spatial molecular signatures of DH lesions to infer malignant potential, offering a foundation for early stratification and intervention strategies.

Several limitations should be noted. Although the space-for-time approach supports comparison of pathological stages in human tissues, it remains inferential and requires validation through longitudinal sampling. In addition, the proposed leader cell model requires functional validation in experimental systems. Finally, the predictive framework for malignant progression remains exploratory and requires further evaluation in large independent cohorts.

In summary, we present a comprehensive spatial multi-omics atlas of human breast cancer evolution and propose that malignant initiation occurs as early as the DH stage. We demonstrate that EGFR⁺ luminal progenitor-like cells, basal barrier disruption, and microenvironmental remodeling jointly drive the initiation of invasion (Figure 7D). Importantly, spatial molecular features predictive of progression are already present in DH lesions, providing a conceptual and translational basis for early risk stratification and interception of breast cancer progression.

## DATA AVAILABILITY

The mass spectrometry proteomics data have been deposited to the ProteomeXchange Consortium (https://proteomecentral.proteomexchange.org) via the iProX partner repository^57,58^ with the dataset identifier PXD079487.

## ACKNOWLEDGMENTS

This work was supported by the Natural Science Foundation of Zhejiang Province (grant no. LHZSD25H160001) and the National Natural Science Foundation of China (grant nos. 82203610, 82302803 and 82300697). We thank Ziqian Wan and the Mass Spectrometry Core Facility of the Central Laboratory, the First Affiliated Hospital, Zhejiang University School of Medicine, for technical support.

## AUTHOR CONTRIBUTIONS

Conceptualization, F.J. and L.W.; methodology, F.J. and L.W.; Investigation, S.J., Z.L., L.J., Y.C., J.S., M.L., D.C., M.Z., Q.F., Q.H., Y.X., J.L., L.Z., Xia.Yang., Y.H., Y.W., and Xuanjing.Ye.; Resources, L.W., L.J., J.S., and Y.L.; Software, Z.L. and M.L.; Data Curation, Y.C., Q.H., J.Z., and Q.M.; Writing-original draft, S.J., F.J., Z.L. and M.L.; Writing-review and editing, all authors; Visualization, Z.L., M.L., X.L., and S.J.; Supervision, F.J., L.W. and J.Z.; Funding acquisition, L.W., F.J., Q.F., and Q.H.;

## EXPERIMENTAL MODEL AND STUDY PARTICIPANT DETAILS

### Patient Selection and Cohorts

#### Multi-stage Samples from Patients with Invasive Ductal Carcinoma of the Breast

##### Inclusion and Exclusion Criteria

Patients diagnosed with invasive ductal carcinoma (IDC) of the breast at the Department of Surgical Oncology, Sir Run Run Shaw Hospital, Zhejiang University School of Medicine, were prospectively enrolled in this study. Inclusion criteria were as follows: (i) age ≥18 years; (ii) histopathologically confirmed invasive ductal carcinoma of the breast; and (iii) provision of written informed consent for participation. Exclusion criteria included: (i) prior history of any malignancy or receipt of antitumor therapies such as radiotherapy or chemotherapy for other diseases; and (ii) vulnerable populations, including individuals with psychiatric disorders, cognitive impairment, critical illness, minors, pregnant women, and illiterate individuals. Withdrawal criteria included: (i) voluntary withdrawal by the participant; or (ii) investigator determination that continued participation was inappropriate. The use of patient tissues and clinical data was approved by the Ethics Committee of Sir Run Run Shaw Hospital, Zhejiang University School of Medicine (Approval No. 2022-497-02).

##### Sample Collection and Preservation

Five to ten breast tissue specimens were collected from surgical resection samples, including tumor tissue, adjacent non-tumorous tissue, and grossly normal breast tissue. Each specimen was divided into two portions. One portion was formalin-fixed and paraffin-embedded (FFPE) for pathological evaluation and single-cell spatial omics analyses. The other portion was snap-frozen in liquid nitrogen and stored at −80°C for multi-omics profiling, including whole-exome sequencing (WES), transcriptomics, proteomics, metabolomics, and 16S microbiome analyses. A unified coding and labeling system was established to comprehensively document sample source, collection time, and storage conditions, thereby ensuring sample traceability. All specimens were processed according to standardized protocols to minimize technical bias.

##### Pathological Evaluation and Sample Selection

FFPE sections were subjected to hematoxylin and eosin (H&E) staining and independently reviewed by three experienced pathologists. Samples were classified as normal breast tissue, usual breast hyperplasia, ductal carcinoma in situ (DCIS), or invasive ductal carcinoma (IDC). Patients harboring at least three distinct pathological stages within the same individual were selected for subsequent analyses.

#### Adjacent Normal Breast Tissue Samples from Patients with Fibroadenoma

##### Inclusion and Exclusion Criteria

Patients clinically diagnosed with breast fibroadenoma at the Department of Surgical Oncology, Sir Run Run Shaw Hospital, Zhejiang University School of Medicine, were enrolled in this study. Inclusion criteria were: (i) age ≥18 years; (ii) clinical diagnosis of breast fibroadenoma; and (iii) provision of written informed consent. Exclusion criteria included: (i) prior history of malignancy or receipt of breast radiotherapy, chemotherapy, or other antitumor treatments; (ii) vulnerable populations, including individuals with psychiatric disorders, cognitive impairment, critical illness, minors, pregnant women, and illiterate individuals; and (iii) pathological diagnosis inconsistent with fibroadenoma. Withdrawal criteria included: (i) voluntary withdrawal by the participant; or (ii) investigator determination that continued participation was unsuitable.

##### Sample Collection and Preservation

One to two grossly normal breast tissue specimens were collected from regions adjacent to the fibroadenoma lesion during surgical resection. Each specimen was divided into two portions. One portion was formalin-fixed and paraffin-embedded for pathological evaluation and single-cell spatial omics analyses. The other portion was snap-frozen in liquid nitrogen and stored at −80°C for multi-omics analyses, including WES, transcriptomics, proteomics, metabolomics, and 16S microbiome profiling.

##### Pathological Evaluation and Sample Selection

FFPE sections were stained with H&E and independently evaluated by three pathologists to confirm tissue identity. Only samples pathologically verified as normal breast tissue were retained for downstream analyses.

##### Cohort for Risk Assessment of Malignant Transformation in Benign Breast Tumors

Patients initially diagnosed and treated at Sir Run Run Shaw Hospital, Zhejiang University School of Medicine, for benign breast tumors-including fibroadenoma and intraductal papilloma-were retrospectively collected. Eligible patients underwent surgical resection for the primary benign lesion and were subsequently rehospitalized at least one year later with a benign or malignant lesion in the ipsilateral breast. Inclusion criteria were: (i) complete pathological diagnoses available for both surgical procedures; (ii) initial diagnosis confirmed as a benign breast lesion; (iii) recurrent lesion located in the ipsilateral breast; and (iv) complete follow-up data available. Exclusion criteria included: (i) prior radiotherapy or chemotherapy; (ii) history of other systemic malignancies; and (iii) inadequate tissue preservation quality.

FFPE tissue blocks from both the initial and recurrent surgeries were retrieved from the Department of Pathology and assembled into tissue microarrays (TMAs). Comprehensive clinical information was systematically collected, including age, family history, imaging characteristics, pathological subtype, tumor size, surgical margin status, hormone receptor status, Ki-67 index, and follow-up outcomes. According to the subsequent ipsilateral lesion, patients were classified into benign-after-benign and malignant-after-benign outcome groups. A unified clinical database with standardized coding management was subsequently established.

## METHOD DETAILS

### Reagents and Resources

Protease inhibitor cocktail, methyl tert-Butyl ether (MTBE; Aladdin, Cat#G2405126), dithiothreitol (DTT; Sigma, Cat#43816), chloroacetamide (CAA; Aladdin, Cat#C106072), trypsin (Bairui, Cat#BELT001), trifluoroacetic acid (TFA; Thermo Fisher Scientific, Cat#85183), acetonitrile (ACN; Aladdin, Cat#A120771-2.5L), Urea (Sigma, Cat#U5378), Polyclonal Rabbit anti-NLGN4Y Antibody (Novus, Cat#NBP2-93145), Polyclonal Rabbit anti-GABRG3 (Novus, Cat#NBP2-30464), Monoclonal Mouse anti-IFNL1 (Novus, Cat# NBP3-41961), Monoclonal Rabbit anti-MGP (Abcam, Cat#ab325749), Monoclonal Rabbit anti-FASLG (Abcam, Cat#ab133619), Polyclonal Rabbit anti-THBS1 (Proteintech, Cat#18304-1-AP), Polyclonal

Rabbit anti-TMSB10 (Abcam, Cat#ab198738), Monoclonal Rabbit anti-FN1 (Abcam, Cat#ab268020), Monoclonal Rabbit anti-TAGLN (CST, Cat#36090T), Monoclonal Mouse anti-MLPH (Thermo Fisher Scientific, Cat#MA5-26992), Monoclonal Mouse anti-IFNE (Novus, Cat#MAB9147-SP), Polyclonal Sheep anti-DISC1 (Novus, Cat#AF6699-SP), Monoclonal Rabbit anti-AZGP1 (Bio-Techne, Cat#NBP3-15426), Monoclonal Mouse anti-KRT8/18 (Abcam, Cat#ab269768), Monoclonal Mouse anti-KRT8/18 (Abcam, Cat#ab17139), Monoclonal Rabbit anti-ACTA2 (Abcam, Cat#ab124964), Monoclonal Rabbit anti-PIP (Thermo Fisher Scientific, Cat# MA5-34862), and protease inhibitor cocktail (Sangon, Cat#C600387) were used in this study.

### Tissue Sample Processing

Breast tissue specimens stored at −80°C were first pulverized under liquid nitrogen. The disrupted tissue powder was transferred into tissue homogenization tubes containing an appropriate amount of fine tissue disruption beads. Subsequently, 56 μL DEPC-treated water supplemented with 2× protease inhibitor cocktail was added, and samples were homogenized using a tissue disruptor for three cycles of 5 s homogenization followed by 5 s rest intervals. An additional 360 μL methanol was then added, followed by one further round of homogenization. After centrifugation, 1200 μL MTBE was added to the homogenate. Samples were vortexed thoroughly and incubated on ice with shaking at 300 rpm for 10 min. Subsequently, 300 μL DEPC-treated water containing 2× protease inhibitor cocktail was added, followed by vortex mixing. After standing on ice for approximately 1 min to allow phase separation, samples were centrifuged at 15,000 × g at 4°C for 10 min. The middle phase was collected for metabolomics analysis, whereas the lower precipitated phase was used for whole-exome sequencing, transcriptomic profiling, and proteomic analyses.

### Whole-Exome Sequencing

#### DNA Extraction and Library Preparation

Genomic DNA was isolated from tissue pellet samples using the MagPure Stool/Soil DNA KF Kit (Magen, Cat#D6356-F-96-SH) according to the manufacturer’s instructions. DNA concentration and purity were assessed using a NanoDrop spectrophotometer (Thermo Fisher Scientific, Wilmington, DE, USA), while DNA integrity was evaluated by 1% agarose gel electrophoresis.

For whole-exome sequencing (WES), genomic DNA libraries were prepared using the Agilent SureSelect Human All Exon V8 kit (Agilent Technologies, Santa Clara, CA, USA) following the manufacturer’s standard workflow. Briefly, genomic DNA was enzymatically fragmented into short inserts and subsequently purified. End repair, A-tailing, and adapter ligation were performed using the reagents provided in the kit. Adapter-ligated fragments were amplified by polymerase chain reaction (PCR) to generate pre-capture libraries. The amplified libraries were then hybridized with exon-targeted capture probes, followed by washing and elution to enrich exonic regions. The enriched libraries were sequenced on the Illumina NovaSeq 6000 platform (Illumina, San Diego, CA, USA) using a 150 bp paired-end sequencing strategy. Whole-exome sequencing and primary bioinformatic analyses were performed by OE Biotech Co., Ltd. (Shanghai, China).

### Whole-Exome Sequencing Data Processing and Analysis

Raw sequencing data were initially generated in FASTQ format. To ensure high-quality downstream analyses, raw reads were processed using fastp (v0.20.0)^59^. Adapter sequences were removed, and low-quality bases within sliding windows with an average Phred quality score below 20 were trimmed. Reads containing ambiguous nucleotides or shorter than 75 bp after filtering were excluded from further analysis.

Clean reads were aligned to the human reference genome (GRCh37.p13) using the Burrows–Wheeler Aligner (BWA) (v0.7.17)^60^. The resulting alignment files were subsequently sorted and indexed using SAMtools (v1.9)^61^. Base quality score recalibration and local realignment around insertion/deletion (INDEL) regions were conducted using the Genome Analysis Toolkit (GATK) (v4.1.9.0)^62^. Single-nucleotide variants (SNVs) and INDELs were identified and annotated using ANNOVAR^63^ with reference to multiple public databases, including RefSeq, the 1000 Genomes Project, the Catalogue of Somatic Mutations in Cancer (COSMIC), and Online Mendelian Inheritance in Man (OMIM). Copy number variations (CNVs) were inferred using CNVkit (v0.9.8)^64^, whereas structural variations (SVs) were detected using LUMPY (v0.2.13)^65^.

### Transcriptome Sequencing

#### RNA Isolation and Library Construction

Total RNA was extracted from tissue pellet samples using the Total RNA Extraction Kit 2.0 Plus (Vazyme, Cat#R411-C3) in accordance with the manufacturer’s instructions. RNA concentration and purity were determined using a NanoDrop 2000 spectrophotometer (Thermo Fisher Scientific, USA), while RNA integrity was evaluated with the Agilent 2100 Bioanalyzer system (Agilent Technologies, Santa Clara, CA, USA).

Sequencing libraries were subsequently generated using the VAHTS Universal V10 RNA-seq Library Prep Kit (Premixed Version) following the manufacturer’s recommended protocol. Transcriptome sequencing and primary bioinformatic analyses were carried out by OE Biotech Co., Ltd. (Shanghai, China).

#### RNA Sequencing and Data Processing

Constructed libraries were sequenced on the Illumina NovaSeq X Plus platform using a 150 bp paired-end sequencing strategy. Approximately 6.9 Gb of raw sequencing data were generated per sample. Raw reads in FASTQ format were initially processed using fastp^59^ to remove adaptor contamination and low-quality reads, yielding approximately 6.8 Gb of high-quality clean reads per sample for downstream analyses.

Clean reads were aligned to the human reference genome using HISAT2^66^. Gene expression abundance was quantified as fragments per kilobase of transcript per million mapped reads (FPKM)^67^, and raw read counts for each gene were calculated using HTSeq-count^68^.

### Proteomics

#### Protein Digestion and Desalting

Protein pellets obtained after tissue processing were resuspended in an appropriate volume of urea lysis buffer and thoroughly homogenized by vortexing. The samples were then subjected to ultrasonication in an ice-water bath for 5 min, followed by centrifugation at 15,000 × g for 15 min at 4°C, and the supernatant was collected.

Protein concentration was determined using a BCA assay. Dithiothreitol (DTT) was added to a final concentration of 5 mM and incubated at 30°C for 30 min for reduction. After cooling to room temperature, chloroacetamide (CAA) was added to a final concentration of 15 mM for alkylation, followed by incubation at room temperature in the dark for 30 min. Equal amounts of protein from each sample, quantified by BCA, were then collected and diluted with 3 volumes of 50 mM Tris-HCl (pH 8.0).

Trypsin was added at a trypsin-to-protein mass ratio of 1:50, and samples were digested at 25°C with shaking at 200 rpm for 12 h. The digestion was quenched by adding trifluoroacetic acid (TFA) to a final concentration of 0.5%, followed by incubation at 4°C for 10 min. Samples were centrifuged at 1,500 × g for 10 min at 4°C, and the supernatant containing peptides was collected.

Peptide desalting was performed using C18 solid-phase extraction cartridges following a sequential washing and elution workflow: two washes with acetonitrile (ACN), three washes with 0.1% TFA, peptide loading, three additional washes with 0.1% TFA, and elution with 50% ACN containing 0.1% TFA. The eluted peptides were vacuum-dried and stored for subsequent mass spectrometry analysis.

#### Liquid Chromatography–Mass Spectrometry (LC–MS/MS)

Proteomic data acquisition and analysis were performed by Hangzhou Supercns Biotech Co., Ltd. (Hangzhou, China). For LC–MS/MS analysis, peptides were separated using a 40-min linear gradient at a flow rate of 400 nL/min on a nanoElute 2 nano-liquid chromatography system, which was directly coupled to a timsTOF HT mass spectrometer (Bruker Daltonics). Separation was performed on a self-packed C18 analytical column (15 cm × 100 µm, 1.9 µm particle size). Mobile phase A consisted of water containing 0.1% formic acid, and mobile phase B consisted of acetonitrile containing 0.1% formic acid.

Eluted peptides were analyzed on a timsTOF HT instrument (Bruker Daltonics) equipped with a capillary electrospray source. The electrospray voltage was set to 1.6 kV. MS data were acquired in DIA parallel accumulation–serial fragmentation (diaPASEF) mode across an m/z range of 300–1500 and an ion mobility range of 1.3 to 0.7 1/K0 [V·s/cm^2^]. DIA windows were set from 344 m/z to 1189 m/z using 13 Th isolation windows, with a ramp time of 50 ms. The collision energy was dynamically ramped according to ion mobility, starting from 20 eV at 0.6 1/K0 [V·s/cm^2^] and increasing to 59 eV at 1.6 1/K0 [V·s/cm^2^].

#### Database Search

Raw LC–MS/MS data were processed using DIA-NN (version 2.2.0)^69^. Peptide identification was performed against the human Swiss-Prot reference proteome downloaded from UniProt on August 21, 2025. The false discovery rate (FDR) for peptide identification was controlled at <1%.

Database search parameters were set as follows: trypsin was selected as the proteolytic enzyme with up to one missed cleavage allowed. Carbamidomethylation of cysteine (C) was set as a fixed modification, while oxidation of methionine (M) and protein N-terminal acetylation were specified as variable modifications. The FDR threshold for peptide-spectrum matches (PSMs) was set at 1% (0.01).

### Metabolomics

#### Sample Preparation

Samples stored at −80°C were thawed at room temperature prior to processing. A 400 μL aliquot of each sample was transferred into a 1.5 mL Eppendorf tube, followed by vacuum drying (lyophilization) to dryness. The residue was vortexed for 10 s. Subsequently, 200 μL of pre-chilled methanol/water mixture (4:1, v/v) was added. Samples were vortexed vigorously for 1 min and subjected to ultrasonication in an ice-water bath for 10 min to enhance metabolite extraction. The mixtures were then incubated at −40°C for 12 h to facilitate protein precipitation and metabolite stabilization. After incubation, samples were centrifuged at 12,000 rpm for 20 min at 4°C. A 120 μL aliquot of the supernatant was transferred into LC-MS vials with glass inserts for subsequent analysis. Quality control (QC) samples were prepared by pooling equal aliquots from all samples to ensure analytical stability and reproducibility across runs.

#### LC–MS/MS Analysis

Metabolomic profiling and data acquisition were performed by Shanghai Luming Biological Technology Co., Ltd. (Shanghai, China). Metabolites were analyzed using an ACQUITY UPLC I-Class Plus system (Waters Corporation, Milford, MA, USA) coupled to a Q Exactive HF mass spectrometer (Thermo Fisher Scientific, Waltham, MA, USA) equipped with a heated electrospray ionization (ESI) source. Data were acquired in both positive and negative ion modes.

Chromatographic separation was performed on an ACQUITY UPLC HSS T3 column (1.8 μm, 2.1 × 100 mm). The mobile phase consisted of (A) water containing 0.1% formic acid (v/v) and (B) acetonitrile containing 0.1% formic acid (v/v). The gradient elution program was set as follows: 0 min, 5% B; 2 min, 5% B; 4 min, 30% B; 8 min, 50% B; 10 min, 80% B; 14 min, 100% B; 15 min, 100% B; 15.1 min, 5% B; and 16 min, 5% B. The flow rate was maintained at 0.35 mL/min, and the column temperature was set at 45°C. During analysis, all samples were kept at 10°C, and the injection volume was 5 μL.

The mass spectrometer was operated over an m/z range of 80–1,200. Full MS scans were acquired at a resolution of 60,000, and HCD MS/MS scans were acquired at a resolution of 15,000. Collision energies were set at 10, 20, and 40 eV. The instrument parameters were as follows: spray voltage, 3,800 V (positive mode) and 3,200 V (negative mode); sheath gas flow rate, 35 arbitrary units; auxiliary gas flow rate, 8 arbitrary units; capillary temperature, 320°C; auxiliary gas heater temperature, 350°C; and S-lens RF level, 50.

#### Data Preprocessing

Raw LC–MS data were processed using Progenesis QI software (v2.3; Nonlinear Dynamics, Newcastle, UK) for baseline correction, peak detection, peak integration, retention time alignment, peak matching, and normalization. The main processing parameters were set as follows: precursor ion tolerance of 5 ppm, product ion tolerance of 10 ppm, and 5% product ion intensity threshold.

Metabolite identification was performed based on accurate mass-to-charge ratio (m/z), MS/MS fragmentation patterns, and isotopic distributions, with reference to multiple databases, including the Human Metabolome Database (HMDB), LIPID MAPS (v2.3), METLIN, and an in-house spectral library.

The resulting feature table was further filtered by removing signals with missing values (ion intensity = 0) in more than 50% of samples within any group. Zero values were replaced with half of the minimum detected value. Metabolites were further filtered based on identification quality; compounds with scores below 36 (out of 60) were excluded. Finally, data matrices from positive and negative ion modes were merged for downstream statistical analyses.

### 16S rRNA Sequencing

#### DNA Extraction and Amplification

Total genomic DNA was extracted using the MagPure Soil DNA LQ Kit (Magen, Cat#D6356-02) according to the manufacturer’s instructions. DNA concentration and integrity were assessed using a NanoDrop 2000 spectrophotometer (Thermo Fisher Scientific, USA) and 1% agarose gel electrophoresis, respectively. Extracted DNA was stored at −20°C until further processing.

The purified DNA was used as a template for PCR amplification of bacterial 16S rRNA genes using barcoded primers and Takara Ex Taq polymerase (Takara). For bacterial community profiling, the V3–V4 hypervariable regions of the 16S rRNA gene were amplified using the universal primers 343F (5′-TACGGRAGGCAGCAG-3′) and 798R (5′-AGGGTATCTAATCCT-3′)^70^.

#### Library Construction and Sequencing

PCR amplicon quality was evaluated by agarose gel electrophoresis. The amplified products were purified using AMPure XP beads (Agencourt) and subjected to a second round of PCR amplification when necessary. After a second purification step with AMPure XP beads, final amplicons were quantified using the Qubit dsDNA HS Assay Kit (Thermo Fisher Scientific, USA). Library concentrations were normalized prior to sequencing.

Sequencing was performed on the Illumina NovaSeq 6000 platform (Illumina Inc., San Diego, CA, USA) using a 250 bp paired-end sequencing strategy, with support from OE Biotech Co., Ltd. (Shanghai, China).

#### Bioinformatic Analysis

Library sequencing and downstream data processing were performed by OE Biotech Co., Ltd. (Shanghai, China). Raw sequencing data were generated in FASTQ format. Paired-end reads were processed using Cutadapt for adapter trimming. Subsequently, reads were filtered for quality control, denoised, merged, and subjected to chimera removal using DADA2^71^ within the QIIME2^72^ pipeline (v2020.11) with default parameters.

The resulting output included representative amplicon sequence variants (ASVs) and an ASV abundance table. Representative sequences for each ASV were selected using the QIIME2 platform. Taxonomic annotation was performed by aligning representative sequences against the SILVA database (v138) using the q2-feature-classifier plugin with default parameters.

### Single-Cell Spatial Transcriptomics (SMI)

#### Sample Preparation for SMI

Single-cell spatial molecular imaging (SMI) was performed by Shanghai Outdo Biotech Co., Ltd. (Shanghai, China). Spatial transcriptomic profiling was conducted using formalin-fixed paraffin-embedded (FFPE) tissue sections according to the CosMx™ SMI Manual Slide Preparation protocol for RNA assays (MAN-10184).

Briefly, FFPE sections were subjected to antigen retrieval at 100°C for 15 min in 1× Target Retrieval Solution, followed by enzymatic digestion with 3 μg/mL Proteinase K in a hybridization oven at 40°C for 30 min. Subsequently, samples were incubated with 0.001% fiducial markers in 2× SSC-T for 5 min at room temperature in the dark to facilitate spatial registration. To assist tissue visualization and region-of-interest (ROI) selection, samples were stained with antibodies against CD298/B2M, pan-cytokeratin (PanCK), CD45, and DAPI.

#### CosMx SMI Instrument Processing

Prepared slides were loaded onto the NanoString CosMx™ SMI instrument for high-plex imaging. A total of 386 fields of view (FOVs) were acquired across the tissue sections to ensure comprehensive coverage and alignment with corresponding hematoxylin and eosin (H&E)–defined regions. Following probe hybridization, unbound probes were thoroughly washed away, and imaging buffer was applied prior to imaging acquisition.

Fluorescent reporter probes containing photocleavable linkers were sequentially activated via UV illumination, followed by stripping with wash buffer. Cyclic imaging and decoding were then performed by the CosMx instrument to read fluorescent barcodes and assign probe identities, thereby generating in situ spatial transcriptomic detection data.

#### Image Processing and Cell Segmentation

Raw imaging data were processed using a customized in-house SMI analysis pipeline, including image registration, feature detection, and spatial localization. Z-stack images of nuclear (DAPI-positive) and membrane-associated staining were used to define cellular boundaries.

Cell segmentation was performed using the machine learning–based Cellpose algorithm to accurately delineate individual cells and assign transcripts to their corresponding cellular and subcellular compartments. Finally, single-cell gene expression profiles were reconstructed by integrating spatial transcript localization with segmented cellular boundaries.

### Single-Cell Spatial Proteomics

#### Generation of CODEX DNA-Conjugated Antibodies

Single-cell spatial proteomic profiling was performed using the CODEX 60-plex system (Akoya Biosciences), and all experimental procedures were conducted by Hangzhou Infinity Biotechnology Co., Ltd. (Hangzhou, China). Prior to conjugation, PhenoCycler DNA barcodes were assigned to each antibody.

For antibody concentration and purification, 50 μg of each antibody was loaded into a 50 kDa molecular weight cutoff (MWCO) centrifugal filter unit. When the initial volume was less than 100 μL, it was adjusted to 100 μL with 1× PBS. The samples were centrifuged at 12,000 × g for 8 min, and the flow-through was discarded. Subsequently, 450 μL of coupling buffer was added. In parallel, PhenoCycler barcode solutions were prepared by adding 10 μL nuclease-free molecular biology-grade water to each lyophilized barcode vial, followed by mixing each reconstituted barcode with 210 μL coupling buffer. The barcode solution was then applied to the top of each filter unit, and antibody–barcode conjugation was carried out by incubating at room temperature for 2 h. After conjugation, the antibody complexes were purified by transferring 5 μL of the reaction mixture into a new 0.2 mL PCR tube and centrifuging at 12,000 × g for 8 min, discarding the supernatant and adding 450 μL purification buffer; this washing step was repeated three times. Finally, the purified PhenoCycler-conjugated antibodies were collected by adding 100 μL antibody storage buffer to each filter unit followed by centrifugation at 3,000 × g for 2 min, yielding a final volume of approximately 120 μL per antibody.

### CODEX FFPE Tissue Staining and Fixation

FFPE tissue sections were baked at 60°C overnight to ensure complete paraffin melting, followed by sequential deparaffinization and rehydration through immersion in xylene for 15 min, 100% ethanol for 10 min and 15 min, 90% ethanol for 15 min, 70% ethanol for 10 min, and finally distilled water for 30 min. Antigen retrieval was performed by immersing slides in 40 mL of 1× sodium citrate buffer or alkaline retrieval buffer in a 50 mL beaker sealed with aluminum foil, followed by high-pressure heating for 30 min and natural cooling to room temperature, after which samples were briefly rinsed in ddH_2_O for 2 min. Slides were then transferred into hydration buffer for 2 min and equilibrated in staining buffer at room temperature for 30 min. During this step, antibody master mixes were prepared by keeping required antibodies on ice, briefly centrifuging prior to use, calculating the required volumes, and combining each antibody with blocking buffer, followed by gentle mixing and storage on ice until application. For staining, slides were placed in a humidified chamber and incubated with 250 μL of antibody mixture applied to one corner of each section, followed by incubation at room temperature for 4 h. After staining, slides were washed twice by sequential incubation in staining buffer for 5 min each and stored appropriately for subsequent processing.

### CODEX Cyclic Imaging and Data Acquisition

For PhenoCycler-Fusion imaging, stained tissue sections were mounted with a flow chamber and loaded into the PhenoCycler-Fusion system for automated cyclic imaging. Reporter molecules were sequentially introduced to the tissue by the instrument and visualized using fluorescence microscopy. Each imaging cycle enabled simultaneous detection of up to three markers across distinct fluorescence channels, followed by gentle isothermal washing to remove fluorophore-linked reporters. By iteratively repeating cycles with different reporter molecules, the full antibody panel was visualized within the same tissue region in a single experiment. At the completion of each run, all images were processed in parallel and exported as QPTIFF files containing full image stacks and associated metadata.

### Multiplex Immunohistochemistry (mIHC)

#### Tissue Deparaffinization, Rehydration, and Antigen Retrieval

FFPE tissue sections were deparaffinized by sequential immersion in xylene twice (10 min per incubation), followed by rehydration through a graded ethanol series (100%, 95%, 85%, and 75%, 5 min each step), and finally rinsed thoroughly with deionized water to remove residual organic solvents. Endogenous peroxidase activity was quenched by incubating sections in 3% hydrogen peroxide (H_2_O_2_) prepared in methanol for 10 min at room temperature, followed by three washes with phosphate-buffered saline (PBS, pH 7.4) for 5 min each.

Antigen retrieval was performed in preheated Tris-EDTA buffer (10 mM Tris base, 1 mM EDTA, pH 9.0) using a constant-temperature water bath at 95–98°C for 25 min. Sections were then allowed to cool naturally to room temperature for 30 min and washed three times with PBS (5 min each) to remove residual retrieval buffer. Non-specific binding was blocked using 5% bovine serum albumin (BSA) in PBS for 30 min at room temperature, and excess blocking solution was gently removed without additional washing.

### Cyclic TSA Fluorescence Staining and Antibody Stripping

Multiplex protein detection was performed through multiple sequential rounds of staining, in which spectrally distinct and non-overlapping fluorescent tyramide substrates were assigned to different target proteins to minimize spectral crosstalk and ensure independent signal resolution. In each staining cycle, tissue sections were incubated overnight at 4°C in a humidified chamber with a validated primary antibody at a pre-optimized working concentration determined by single-plex IHC titration.

After primary antibody incubation, sections were washed three times with PBS (5 min each) to remove unbound antibodies, followed by incubation with horseradish peroxidase (HRP)-conjugated polymer secondary antibodies matched to the host species of the primary antibody for 30 min at room temperature. After additional PBS washes (3 × 5 min), fluorescent tyramide signal amplification (TSA) reagents (diluted 1:100 in TSA buffer according to manufacturer recommendations) were applied for 10 min at room temperature. During this step, HRP catalyzes covalent deposition of activated fluorescent tyramide onto tyrosine residues proximal to the antigen, generating stable fluorescence signals that are resistant to subsequent stripping procedures.

Following each staining cycle, antibody complexes were removed by incubating sections in preheated citrate buffer (10 mM citric acid, 0.05% Tween-20, pH 6.0) at 98°C for 15 min. This stripping step efficiently eliminated bound primary and secondary antibodies while preserving covalently deposited fluorescent signals, thereby preventing signal carryover between cycles. Sections were then cooled to room temperature, washed three times with PBS (5 min each), and subjected to the next staining cycle using a different target antibody and spectrally distinct TSA fluorophore. After completion of all staining cycles, nuclei were counterstained with 4′,6-diamidino-2-phenylindole (DAPI, 1 μg/mL in PBS) for 5 min at room temperature, followed by three final PBS washes.

### Slide Mounting, Multispectral Imaging, and Analysis

Stained sections were mounted using an anti-fade fluorescence mounting medium to minimize photobleaching and ensure signal stability during imaging and long-term storage, and coverslips were sealed with clear nail polish to preserve tissue integrity. Whole-slide multispectral imaging was performed using an automated quantitative pathology imaging system equipped with narrow-band excitation and emission filters corresponding to each fluorescent channel and DAPI.

Spectral unmixing was conducted using dedicated multispectral image analysis software to separate true signal from tissue autofluorescence and eliminate residual spectral overlap. This enabled high-resolution single-cell–level quantification and precise spatial localization of multiplex protein expression across tissue sections.

## QUANTIFICATION AND STATISTICAL ANALYSIS

### Transcriptomic Data Analysis

Genes detected with expression > 0 in at least one sample were defined as qualitatively detected genes, whereas genes expressed in ≥50% of samples were retained as quantitatively detected genes. For downstream analysis, the selected quantitative gene expression matrix was transformed using log_2_(x + 1) to stabilize variance and accommodate zero values. Subsequently, Pareto scaling was applied using the formula (x − mean(x)) / √sd x to reduce the influence of highly abundant genes.

Differential expression analysis was performed using the DESeq2 package (v1.40.2)^73^. Prior to analysis, sample mapping tables were used to extract corresponding count matrices for each comparison group. Two experimental designs were considered: for unpaired comparisons, a model formula of ∼ group was applied; for paired designs (samples derived from the same patient), a ∼ patient + group model was used to account for inter-individual variability. During DESeq2 processing, if the default model fitting failed, local regression and mean-based fitting were sequentially attempted as fallback strategies. Differentially expressed genes (DEGs) were defined as those with *P* < 0.05 and |log_2_FC| > 1. Based on expression direction, DEGs were classified into upregulated genes (log_2_FC > 1) and downregulated genes (log_2_FC < −1).

### Proteomic Data Analysis

Proteins detected with abundance > 0 in at least one sample were defined as qualitatively detected proteins, while proteins detected in ≥50% of samples were retained as quantitatively detected proteins. The quantitative protein matrix was log_2_(x + 1) transformed to stabilize variance and handle missing values. Median normalization was applied to correct for systematic bias between samples, where the expression value of each sample was subtracted by the median of all protein expression values within that sample, aligning the medians of all samples to zero, thereby reducing the impact of systematic technical variations.

Differential protein expression analysis was performed using the limma package (v3.56.2)^74^. Prior to analysis, median normalization was applied to the protein expression matrix to ensure comparability across samples. Two experimental designs were considered: for unpaired comparisons, a design matrix of ∼0 + group was used to estimate group-specific coefficients; for paired designs (samples derived from the same patient), a ∼0 + group + patient model was applied, and intra-patient correlation was estimated using the duplicateCorrelation function to account for repeated measures. After linear model fitting, empirical Bayes moderation was applied using the eBayes function, shrinking protein-specific variances toward a pooled prior estimate to improve stability under small sample sizes. Contrast matrices were constructed using the makeContrasts function according to the study design, and moderated t-statistics were computed. Differentially expressed proteins (DEPs) were defined as those with *P* < 0.05 and |log_2_FC| > 1, and were categorized into upregulated proteins (log_2_FC > 1) and downregulated proteins (log_2_FC < −1).

### Metabolomic Data Analysis

Differential metabolite identification was performed using orthogonal partial least squares discriminant analysis (OPLS-DA)^75^. Prior to modeling, missing values in the normalized metabolite matrix were imputed using median-based replacement. OPLS-DA was implemented using the ropls package (v1.32.0), with one predictive component and one orthogonal component. The number of cross-validations was set to min (7, sample size − 1), and permutation testing was performed with 1,000 iterations. If OPLS-DA modeling failed, a partial least squares discriminant analysis (PLS-DA) model was automatically applied as a fallback strategy. Model performance was evaluated using R^2^Y (explained variance of group separation) and Q^2^ (predictive ability). Variable importance in projection (VIP) scores were used to identify metabolites contributing to group discrimination (VIP > 1).

Statistical significance was further assessed using Student’s t-test between experimental and control groups, followed by multiple testing correction using the Benjamini–Hochberg (BH) procedure to control false discovery rate. Differential metabolites were defined as those meeting VIP > 1, *P* < 0.05, and |log_2_FC| > 1, and were classified into upregulated (log_2_FC > 1) and downregulated metabolites (log_2_FC < −1).

### 16S Microbiome Data Analysis

Differential abundance analysis was performed using the MaAsLin2 package (v1.15.1)^76^. Prior to analysis, ASV (amplicon sequence variant) abundance data were normalized to relative abundance and log_2_(x + 1) transformed for modeling. Two analytical designs were applied: for unpaired comparisons, the model Feature ∼ Group + Age + BMI was used, with age and BMI included as covariates; for paired designs (samples derived from the same patient), a mixed-effects model Feature ∼ Group + Age + BMI + (1|Patient_ID) was used to account for intra-individual variability.

MaAsLin2 was run using a linear model (LM) framework without additional normalization or transformation. Minimum abundance and minimum prevalence thresholds were set to 0. Multiple testing correction was performed using the Benjamini–Hochberg (BH) method, with a significance threshold of *P* < 0.05. Differential ASVs were defined as those with adjusted *P* < 0.05 and |log_2_FC| > log_2_(1.5), and were classified into increased (log_2_FC > log_2_(1.5)) and decreased (log_2_FC < −log_2_(1.5)) groups.

### Gene Set Enrichment Analysis (GSEA)

GSEA was performed using the gseGO and gseKEGG functions in the clusterProfiler package (v4.8.3)85. Genes were ranked based on their mutation frequency in each comparison group, with higher mutation frequency ranked higher. GSEA for Gene Ontology terms was performed using the gseGO function separately for biological process (BP), cellular component (CC), and molecular function (MF) sub-ontologies. KEGG pathway GSEA was performed using the gseKEGG function. Parameter settings were as follows: P value threshold < 0.05, number of permutations (nPermSimple) = 1000, minimum gene set size (minGSSize) = 5, maximum gene set size (maxGSSize) = 800.

### Spatial Proteomics Image Processing and Cell Segmentation

Spatial proteomics images were analyzed using QuPath software^77^. Nuclear segmentation and whole-cell boundary delineation were performed based on the StarDist2D deep learning algorithm^78^. The DAPI channel was used as the primary input for nuclear detection. Standardized segmentation parameters were applied across all images, including percentile normalization (1%–99%), a cell detection probability threshold of 0.5, a spatial resolution of 0.5 μm per pixel, and a cytoplasmic expansion distance of 5 μm from the nucleus. Batch-wise standardized cell detection was performed across all samples. After quality control and integration, a total of 622,760 single cells were retained for downstream spatial proteomic classification and quantitative analyses.

### Spatial Proteomics Data Preprocessing, Dimensionality Reduction, and Clustering

Downstream analyses were conducted in R (v4.4.2) using the Seurat package (v5.4.0)^79^. Raw spatial proteomics data, sample metadata, and subcellular protein localization annotations were integrated to construct a unified dataset. Cellular morphological features and mean protein expression matrices were extracted and merged with metadata.

Comprehensive quality control was performed, including distributions of cell counts per sample, total protein signal intensity per cell, and single-marker expression profiles. Cells were filtered using the following thresholds: DAPI intensity between 10 and 250, total protein expression > 0, and cell area > 0. After filtering, 606,467 high-quality cells were retained.

Protein expression matrices were integrated with spatial coordinates, morphological features, and sample annotations to construct Seurat objects. Data were normalized using the LogNormalize method, and sample identity was used as the batch variable for downstream correction.

All 60 protein features were scaled prior to principal component analysis (PCA). The top 30 principal components were extracted, and batch effects across patients were corrected using the Harmony algorithm^80^. Based on elbow plot inspection, the first 10 Harmony dimensions were selected for downstream analysis. A shared nearest-neighbor graph was constructed from the corrected embedding.

To ensure reproducibility, a fixed random seed (42) was used throughout. Cell clustering was performed using the Leiden algorithm^81^ at a resolution of 0.5, resulting in 12 clusters visualized by UMAP^82^. Based on protein expression profiles, clusters were annotated into five major cellular compartments, and poorly defined clusters were excluded. Subclustering was then performed within each major lineage at a resolution of 0.25 using the first 10 PCA and 10 Harmony dimensions. Clusters with low abundance or ambiguous identity were removed during iterative refinement. After final harmonized re-clustering, 521,235 cells were retained. UMAP embeddings and feature protein heatmaps were generated to support cluster annotation.

### Spatial Transcriptomics Data Preprocessing and Clustering

Spatial transcriptomic data were generated using the CosMx 6K platform^83^, comprising 354,282 cells. Quality control included evaluation of per-field-of-view (perFOV) and per-sample distributions, transcript and gene counts per cell, negative control probe counts, and average transcript/gene distributions per FOV.

Cells were filtered using the following criteria: nCount_RNA > 50, negative probe proportion < 5%, and exclusion of cells exceeding the 99th percentile in area. After removal of segmentation outliers, 334,025 high-quality cells were retained.

Data normalization and PCA were performed using scPearsonPCA, followed by batch correction using Harmony. Unsupervised clustering was performed using Seurat FindClusters (resolution = 0.8). Clusters lacking specific marker gene signatures were considered technical artifacts and removed, resulting in 239,599 high-confidence cells. Major cell types were annotated based on canonical marker genes.

Following major annotation, refined subclustering was performed. Prior to subclustering, gene expression overlap was evaluated using the overlap_ratio_metric function in the smiDE package^84^ to remove contaminated or poorly segmented cells. For immune lineages (myeloid, T/NK/B cells), supervised annotation was performed using HieraType^85^, which constructs meta-gene scores based on predefined marker sets and assigns cell identities through a probabilistic framework robust to sparsity and noise.

Before visualization, cluster-level fold-change statistics (log fold change, mean expression, and detection rate) were computed using clusterwise_foldchange_metrics. UMAP embeddings and dot plots of marker genes were generated to visualize cluster structure and expression signatures.

### Cell Density Gradient Analysis

Spatial cell density was quantified using single-cell spatial coordinates from both spatial transcriptomics and proteomics datasets. Tissue areas were estimated using a concave hull algorithm^86^. Spatial transcriptomics coordinates were converted using a scaling factor of 0.12028 μm per pixel, whereas spatial proteomics data were directly analyzed in native micrometer coordinates.

Cell density was defined as the number of cells per square millimeter of tissue. Boxplots were generated to compare density distributions across disease stages.

### Cell Morphology Analysis

Cell morphological features, including area, minor axis length, major axis length, and eccentricity, were extracted from both spatial transcriptomic and proteomic datasets. After integration with cell annotations, mean values (two decimal precision) were calculated across cell types, subclusters, and disease stages.

### Spatial Mapping of Cellular Populations

Spatial coordinates and annotations were integrated to visualize global and epithelial subpopulation distributions across disease stages. Spatial proteomics data were rendered using GeoJSON-based geometry^87^, while spatial transcriptomic data were visualized using CosMx polygon coordinates. Pixel-to-micron conversion (0.12028) was applied to transcriptomic data, whereas proteomic data were analyzed in native micron-scale coordinates. All spatial plots included micron-scale rulers and were generated per sample/FOV^88^. Cells labeled as “Unknown” were excluded.

### Basal Gap Cell Analysis

Basal gap cells were defined as epithelial cells located in regions of disrupted basal layer continuity, characterized by loss of mature basal epithelial cells in mammary duct structures. These cells were manually annotated based on spatial maps integrating transcriptomic and proteomic data with ductal morphology. A total of 5,100 and 5,683 basal gap cells were identified in spatial transcriptomics and proteomics datasets, respectively.

For each sample, the proportion and spatial density of basal gap cells relative to total basal epithelial cells were calculated.

### Mammary Duct Analysis

The number of mammary ducts was quantified from H&E images, spatial transcriptomic data, and spatial proteomic images across disease stages. Boxplots were generated at the sample level to assess distributional differences and trends across progression stages.

### Differential Gene Expression Analysis of Basal Cells

Differential gene expression analysis across different stages of breast cancer progression was performed on the mBasalEpi subpopulation identified from spatial single-cell transcriptomic data using a pseudobulk strategy coupled with the DESeq2 framework. Seven pairwise comparison groups were established according to the sequential progression of pathological stages. Comparisons between BN and MN were conducted using an unpaired design, whereas the remaining six comparisons employed a paired design.

Prior to analysis, quality-control criteria were applied, requiring at least 10 cells in the target cell subpopulation and a minimum of two biological samples per group. Raw RNA count matrices were aggregated using the AggregateExpression function in Seurat^89^, with sample identifiers specified as grouping variables. Gene expression counts from all cells belonging to the same cell subpopulation within each sample were summed to generate a pseudobulk count matrix, with genes represented as rows and samples as columns. Corresponding metadata, including patient identifiers, sample identifiers, and pathological stage annotations, were extracted from the Seurat object.

For each comparison, only samples belonging to the two pathological stages under investigation were retained. In unpaired analyses, all eligible samples were included. In paired analyses, only patients with matched samples from both pathological stages were retained. Pathological stage was encoded as a factor variable, with the reference group specified as the first level. For unpaired comparisons, a DESeq2 generalized linear model incorporating pathological stage as the sole explanatory variable was constructed. For paired comparisons, both pathological stage and patient identity were included as explanatory variables.

Differential expression analysis was performed using the DESeqDataSetFromMatrix function in DESeq2. Size factors were estimated using the median-of-ratios method to correct for differences in sequencing depth and RNA composition among samples. Briefly, geometric mean expression values were first calculated for each gene across all samples to construct a virtual reference sample. For each sample, the ratio between the raw pseudobulk count and the corresponding gene-specific reference value was then computed, excluding genes with zero counts across all samples. The median ratio across all valid genes was subsequently used as the size factor for that sample. Normalized expression values were obtained by dividing raw counts by the corresponding size factors. This normalization strategy minimizes the influence of a small number of highly differentially expressed genes on library size estimation, thereby improving the robustness of statistical inference.

Following normalization, lowly expressed genes were filtered, gene-wise dispersions were estimated, negative binomial generalized linear models were fitted, and Wald tests were performed. Differential expression results were extracted using the DESeq2 results function and compiled into a standardized data frame containing gene name, cell subpopulation, comparison group, log_2_FC, and *P* value. Genes with |log_2_FC| > log_2_(1.5) and adjusted *P* value < 0.05 were considered significantly differentially expressed.

### Differential Expression Analysis of Basal Gap Cells

Basal gap cells were extracted from both spatial datasets and stratified by disease stage. Cell type composition within basal gap regions was quantified and visualized using stacked bar plots.

In spatial transcriptomics, mBasalEpi was defined as mature basal epithelial cells; GATA3mLumEpi, PIPmLumEpi, NPY1RmLumEpi, IFI6mLumEpi, EGFRpLumEpi, PTNEpi, ICAM1Epi, and CRYABEpi were defined as luminal epithelial cells. In spatial proteomics, basalEpi was defined as mature basal cells; basal-likepLumEpi, CD11cmLumEpi, GZMBmLumEpi, iNOSmLumEpi, LAG3mLumEpi, and pLumEpi were defined as luminal populations.

Within-sample paired differential expression analyses were performed for basal gap cells. Two comparison schemes were evaluated: (1) basal gap cells versus basal cells within the same sample, and (2) basal gap cells versus luminal cells within the same sample. Independent analyses were conducted both on the integrated dataset comprising all pathological stages and on datasets stratified by individual pathological stages.

A pseudobulk paired-analysis strategy was adopted. First, all cells belonging to the target cell types were extracted from the original Seurat object. For each sample, the presence of the relevant cell types was assessed, and only samples containing both cell populations under comparison were retained. Samples containing only one of the two cell types were excluded to ensure valid paired comparisons. Subsequently, raw expression matrices were aggregated using the AggregateExpression function in Seurat according to both sample identity and cell type. Expression counts from all cells belonging to the same cell type within each sample were summed, generating a pseudobulk expression matrix in which rows corresponded to genes (or proteins) and columns corresponded to sample–cell-type combinations.

Sample metadata were reconstructed from pseudobulk matrix column names to generate a pairing factor representing sample identity and a grouping factor representing cell type. The grouping factor was ordered such that the control group preceded the experimental group. The aggregated count matrix was converted into a DGEList object using the DGEList function in edgeR. Normalization factors were calculated using the trimmed mean of M-values (TMM) method implemented in calcNormFactors to correct for compositional biases arising from differences in sequencing depth and the presence of highly expressed genes.

Based on TMM normalization, the voom function in the limma package was applied for transformation and secondary normalization. Specifically, discrete integer count data were converted into continuous log-counts per million [log(CPM)] values using effective library sizes and a small pseudocount. Quantile normalization was simultaneously incorporated within the voom procedure to align expression distributions across samples and reduce global systematic technical variation. The voom algorithm further modeled the mean–variance relationship of log(CPM) values using a locally weighted scatterplot smoothing (LOWESS) curve and estimated precision weights for each observation, thereby reducing the influence of lowly expressed and highly variable genes in downstream analyses.

A linear model design matrix incorporating both paired-sample effects and cell-type effects was constructed using the model.matrix function in limma. Weighted linear models were fitted using lmFit, followed by empirical Bayes moderation with eBayes^90^. Multiple testing correction was performed using the Benjamini–Hochberg procedure to control the false discovery rate (FDR). Differential expression results were extracted using topTable and organized into a standardized data frame containing gene name, cell subpopulation, comparison group, log_2_FC, *P* value, and FDR value. Genes or proteins satisfying |log_2_FC| > log_2_(1.5) and FDR < 0.05 were considered significantly differentially expressed.

### Cell-Cell Interaction Analysis

Spatial cell–cell interactions were inferred based on nearest-neighbor Euclidean distances derived from spatial transcriptomic coordinates. Analyses were performed independently for each disease stage using Python (v3.12.13), with stLearn (v1.1.1) and Scanpy (v1.11.5)^91,92^.

Cell coordinates were standardized and processed using stLearn, followed by integration with a curated human ligand–receptor database. A spatial interaction distance threshold of 250 coordinate units and a minimum cell count of 2 were applied. Ligand–receptor interactions were evaluated using 1,000 permutation tests, with FDR-BH correction at the spot level (FDR < 0.05). Cell–cell communication was further assessed using group-level aggregation with identical permutation settings.

For each disease stage, global interaction heatmaps between cell subpopulations were generated. Bar plots were used to visualize the top 20 ligand–receptor pairs ranked by interaction strength between selected cell subpopulations. For specific cell subpopulations, differential expression analysis was performed following the Differential Expression Analysis of Basal Gap Cells workflow, and genes involved in key ligand–receptor pairs were visualized. Spatial interaction networks were constructed by integrating polygon coordinates, neighborhood architecture, and gene expression data, with directional arrows indicating bidirectional ligand-receptor signaling.

### Spatial Niche Analysis

Spatial niche identification was performed independently on spatial transcriptomic and proteomic datasets using Python (v3.11.15). Key tools included Scanpy (v1.11.5), Squidpy (v1.8.1), CellCharter (v0.3.7), and scvi-tools (v1.4.2)^93–95^.

For spatial transcriptomics, scVI was used for dimensionality reduction with batch labels defined by sample identity. For spatial proteomics, precomputed UMAP embeddings were used as initial features with additional sample-level scaling correction. Spatial neighbor graphs were constructed using Delaunay triangulation after removal of long-distance edges. CellCharter was applied to aggregate three layers of spatial neighborhood information into integrated feature representations.

Gaussian mixture modeling (GMM) was used for clustering after parameter optimization, resulting in eight spatial niches in both modalities. Integrated results were visualized in R.

### mIHC Data Analysis

#### Image Segmentation and Single-Cell Protein Expression Quantification

Multiplex immunohistochemistry images were processed using QuPath software. Cell nuclei were detected based on the DAPI fluorescence channel, and whole-cell boundaries were delineated via the StarDist2D deep learning algorithm. Uniform segmentation parameters were applied across all tissue microarray (TMA) images, including 1%–99% percentile normalization, a cell detection probability threshold of 0.5, a spatial resolution of 0.5 μm per pixel, and a cytoplasmic expansion distance of 5 μm from the nuclear boundary. Standardized batch-wise cell detection was performed across all samples.

### Quality Control and Cell Subset Definition

All downstream analyses were performed in R (v4.4.2) using the Seurat package (v5.4.0). Single-cell morphological features and mean fluorescence intensity measurements exported from QuPath were imported and integrated with sample metadata for subsequent processing. Quality control filtering was applied to retain cells with DAPI fluorescence intensity ranging from 5 to 250, total protein expression greater than 0, and cell area greater than 0, to exclude low-quality cells, dead cells, and segmentation artifacts. αSMA-positive cells were defined as cells with αSMA expression ranking in the top 10% of the total cell population and concurrent Cytokeratin 8/18 expression outside the top 10% threshold. Cytokeratin 8/18-positive cells were defined using the corresponding inverse threshold criteria.

### Differential Protein Expression Analysis and Statistical Testing

Two independent TMA cohorts were analyzed with predefined paired and unpaired comparison designs. In the discovery cohort, seven pairwise comparisons were established. The comparison between benign nevus (BN) and malignant nevus (MN) was analyzed using an unpaired design, while the remaining six comparisons (MN vs. DH, MN vs. DCIS, MN vs. IDC, DH vs. DCIS, DH vs. IDC, and DCIS vs. IDC) were analyzed using paired designs based on matched patient samples. In the validation cohort, four disease groups were included: first benign (BF), second benign (BS), first malignant (MF), and second malignant (MS). Unpaired tests were applied for comparisons between BF and MF as well as BS and MS. Paired tests were used for comparisons of longitudinal samples obtained from the same patient at two different time points (BF vs. BS, MF vs. MS).

### Differential expression analysis of mean fluorescence intensity

For each of the three cell datasets (all cells, αSMA-positive cell subset, and Cytokeratin 8/18-positive cell subset), pseudo-bulk mean fluorescence intensity values were calculated per sample by averaging single-cell expression levels, and used for intergroup differential expression analyses. Differential expression analyses were performed using the limma framework. For each comparison, protein expression matrices were fitted with linear models. For paired designs, patient identity was incorporated as a blocking factor in the design matrix to control for inter-individual variation. Statistical significance was assessed using empirical Bayes moderation. Results were summarized as log_2_ fold change (log_2_FC) and *P* values.

### Paired expression comparison between **α**SMA-positive and Cytokeratin 8/18-positive cells

For each disease stage, protein expression levels were compared between αSMA-positive and Cytokeratin 8/18-positive cells derived from the same tissue sample. Only samples containing both cell subsets were retained for analysis. Paired difference testing was performed using the limma framework. Expression differences between the two matched cell subsets from each sample were modeled as the response variable, and statistical significance was evaluated via empirical Bayes moderation. Results were reported as log_2_FC and *P* values.

### Target protein positivity rate analysis in cell subsets

Protein positivity was uniformly defined as expression level above the 90th percentile threshold of the whole cell population (i.e., top 10% high expression). Within the αSMA-positive cell subset (αSMA high expression with concurrent low Cytokeratin 8/18 expression) and Cytokeratin 8/18-positive cell subset respectively, the proportion of cells positive for each target protein was calculated per sample. Intergroup comparisons of positivity rates were performed following the corresponding study design, with unpaired tests for cross-group comparisons and paired tests for matched sample comparisons. Results were presented as percentage ratios and *P* values. Protein expression correlation analysis Spearman correlation coefficients between lineage markers (SMA / Cytokeratin 8/18) and each target protein were calculated at the single-cell level.

---

**Figure S1.**
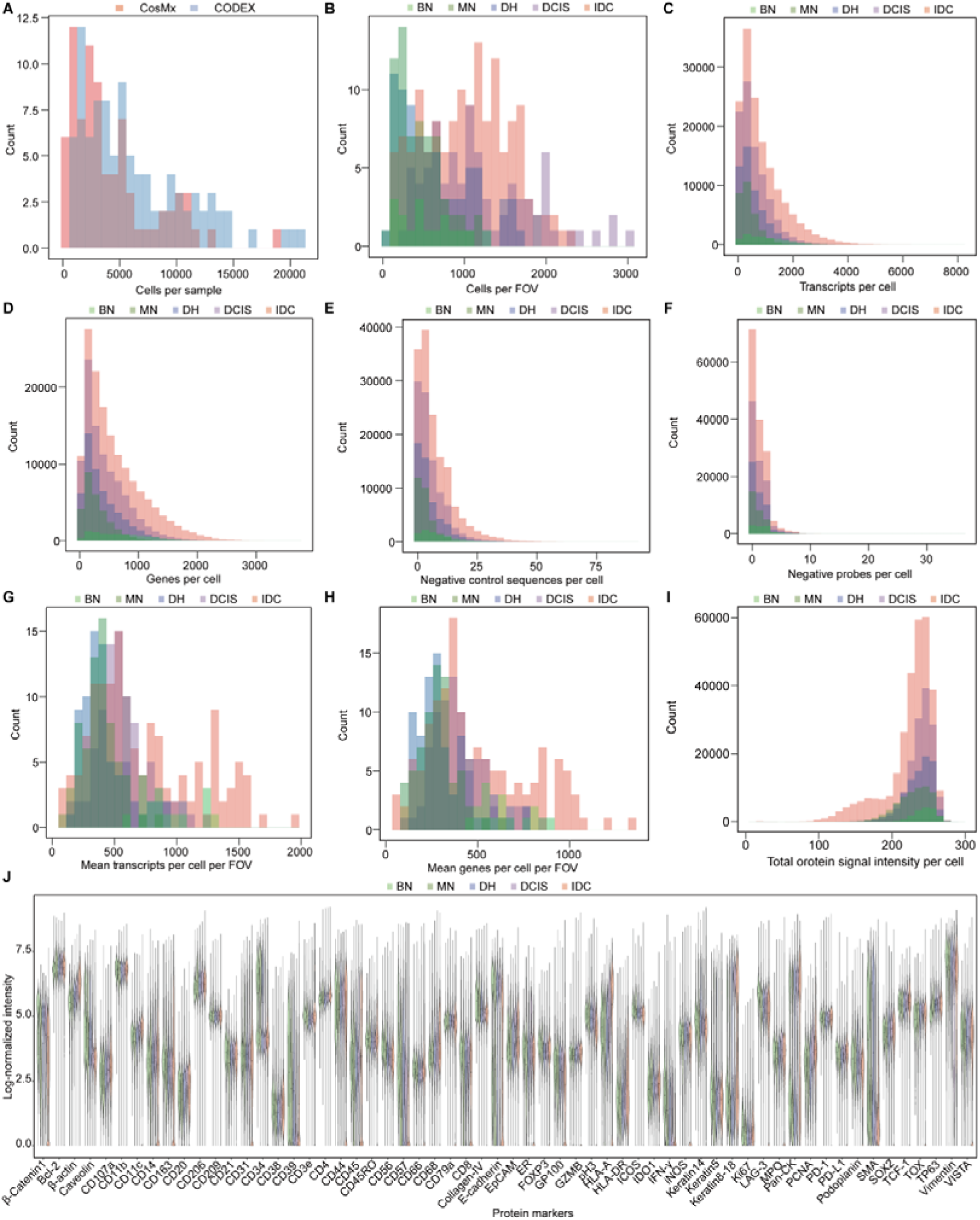
Quality control metrics for CosMx spatial transcriptomics (ST) and CODEX spatial proteomics (SP) data during breast cancer progression, related to figure 2. (A) Stacked density histogram showing the distribution of cells per sample for CosMx (orange) and CODEX (blue) data. (B) Stacked density histogram showing the distribution of cells per field of view (FOV) across BN, MN, DH, DCIS and IDC stages in CosMx data. (C) Stacked density histogram showing the distribution of transcripts per cell across disease stages in CosMx data. (D) Stacked density histogram showing the distribution of genes per cell across disease stages in CosMx data. (E) Stacked density histogram showing the distribution of negative control sequences per cell across disease stages in CosMx data. (F) Stacked density histogram showing the distribution of negative probes per cell across disease stages in CosMx data. (G) Stacked density histogram showing the distribution of mean transcripts per cell per FOV across disease stages in CosMx data. (H) Stacked density histogram showing the distribution of mean genes per cell per FOV across disease stages in CosMx data. (I) Stacked density histogram showing the distribution of total protein signal intensity per cell across disease stages in CODEX data. (J) Violin plot showing the log-normalized expression intensity distribution of all protein markers across disease stages in CODEX data. BN, normal from benign patient; MN, normal from malignant patient; DH, intraductal hyperplasia; DCIS, ductal carcinoma in situ; IDC, invasive ductal carcinoma; FOV, field of view.

**Figure S2.**
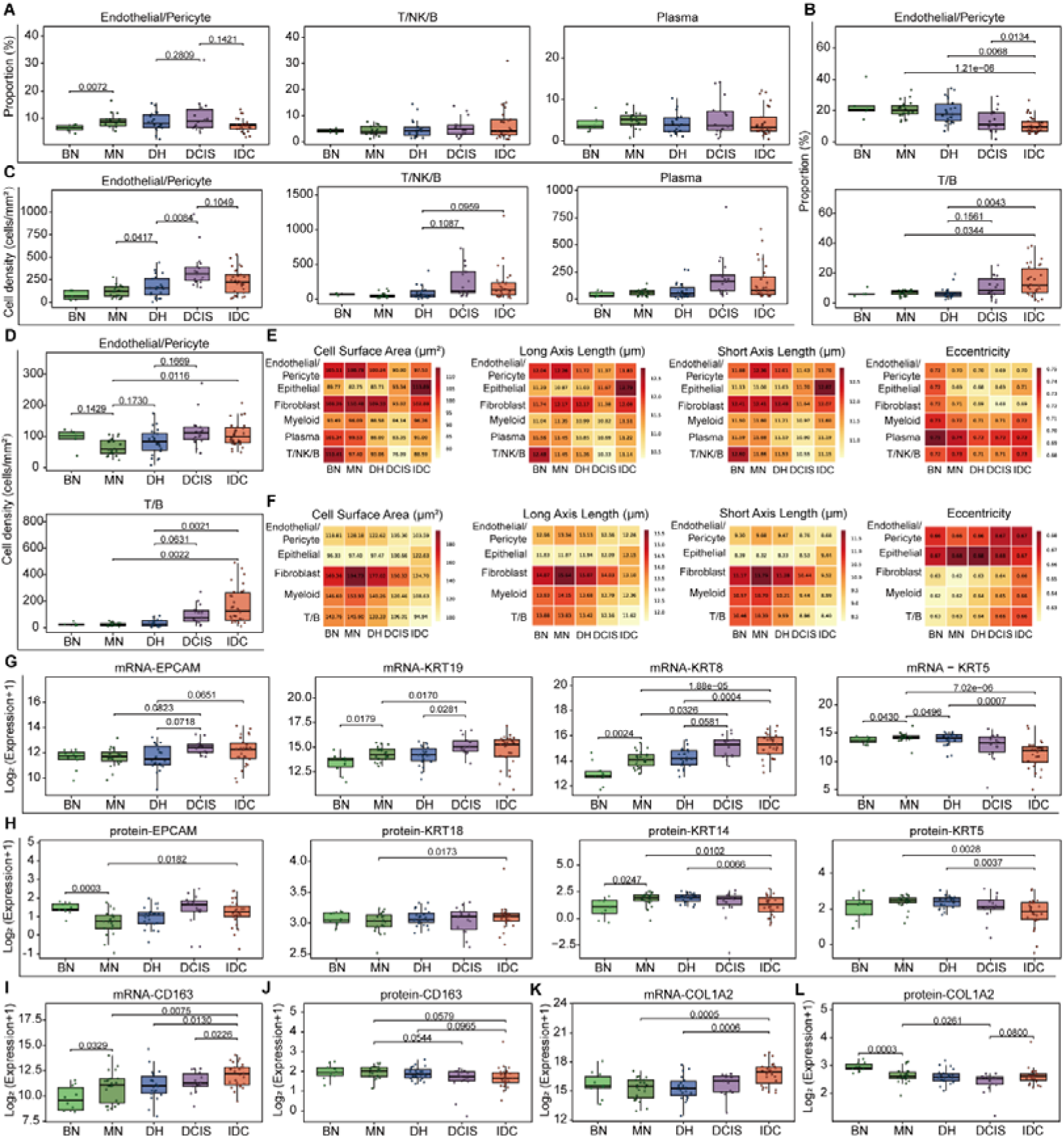
Dynamic changes in spatial cellular composition during breast cancer progression, related to figure 2. (A) Box-and-scatter plots depicting the proportion of Endothelial/Pericyte, T/NK/B and Plasma cells across BN, MN, DH, DCIS and IDC stages in CosMx data. (B) Box-and-scatter plots depicting the proportion of Endothelial/Pericyte and T/B cells across disease stages in CODEX data. (C) Box-and-scatter plots showing cell density (cells/mm^2^) of Endothelial/Pericyte, T/NK/B and Plasma cells across disease stages in CosMx data. (D) Box-and-scatter plots showing cell density (cells/mm^2^) of Endothelial/Pericyte and T/B cells across disease stages in CODEX data. (E) Heatmaps of morphological metrics (cell surface area, long axis length, short axis length, eccentricity) for major cell types across disease stages in CosMx data. (F) Heatmaps of morphological metrics (cell surface area, long axis length, short axis length, eccentricity) for major cell types across disease stages in CODEX data. (G-L) Boxplots showing expression of epithelial markers (EPCAM, KRT19, KRT8, KRT5) in bulk transcriptomics (G) and bulk proteomics (H); myeloid marker (CD163) in bulk transcriptomics (I) and bulk proteomics (J); fibroblast marker (COL1A2) in bulk transcriptomics (K) and bulk proteomics (L). Statistical analysis: Seven pairwise stage comparisons were conducted. Independent-sample tests were used for MN vs BN, whereas paired tests were applied to the other six comparisons using only matched patient samples (DH vs MN, DCIS vs DH, IDC vs DCIS, DCIS vs MN, IDC vs MN, IDC vs DH). All tests were performed using the t test for A, B, C, D, G, H, I, J, K, L. Comparisons without indicated *P* values were not statistically significant. *P* values are presented to four significant digits; *P <* 0.0001 is indicated in scientific notation. BN, normal from benign patient; MN, normal from malignant patient; DH, intraductal hyperplasia; DCIS, ductal carcinoma in situ; IDC, invasive ductal carcinoma.

**Figure S3.**
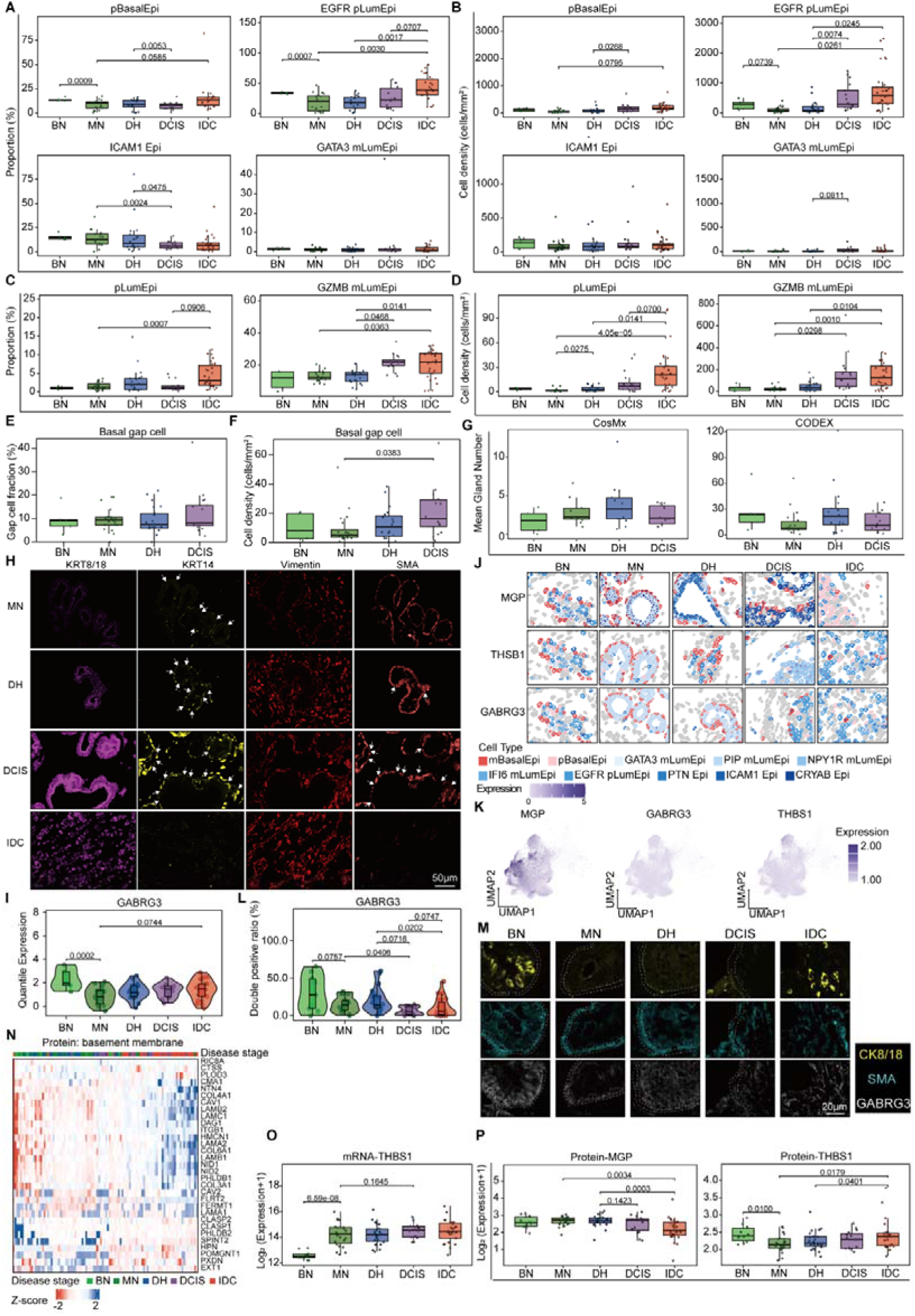
Early basal cell loss and barrier disruption during breast cancer progression, related to figure 3. (A) Box-and-scatter plots depicting the proportion of pBasalEpi, EGFR pLumEpi, ICAM1 Epi and GATA3 mLumEpi subclusters across BN, MN, DH, DCIS and IDC stages in CosMx data. (B) Box-and-scatter plots showing cell density (cells/mm^2^) of pBasalEpi, EGFR pLumEpi, ICAM1 Epi and GATA3 mLumEpi subclusters across disease stages in CosMx data. (C) Box-and-scatter plots depicting the proportion of pLumEpi and GZMB mLumEpi subclusters across disease stages in CODEX data. (D) Box-and-scatter plots showing cell density (cells/mm^2^) of pLumEpi and GZMB mLumEpi subclusters across disease stages in CODEX data. (E) Box-and-scatter plots depicting the proportion of basal gap cells across disease stages in CODEX data. (F) Box-and-scatter plots showing cell density (cells/mm^2^) of basal gap cells across disease stages in CODEX data. (G) Box-and-scatter plots showing the mean gland number across disease stages in CosMx (left) and CODEX (right) data. (H) Representative immunofluorescence images of CODEX data across MN, DH, DCIS and IDC stages, showing the expression of basement membrane-related markers: KRT8/18, KRT14, Vimentin and SMA. Scale bar, 50 μm. White arrows indicate basal gap cells. (I) Violin plots showing the expression distribution of GABRG3 in mBasalEpi subcluster across disease stages. (J) Representative spatial expression maps showing the expression patterns of mBasalEpi differentially expressed genes (DEGs) across pairwise disease stage comparisons, with cell type indicated by border color and gene expression level represented by fill intensity. (K) UMAP projections showing the expression distribution of representative DEGs in epithelial subclusters from CosMx ST data. (L) Violin plots showing the double positive ratio of GABRG3 in SMA+ cell across disease stages. (M) Representative immunofluorescence images of mIHC across MN, DH, DCIS and IDC stages, showing the expression of KRT18/8, SMA, and GABRG3. Scale bar, 20 μm. Two white dashed lines delineate the basal barrier. (N) Heatmap showing the z-score normalized expression levels of basement membrane-associated proteins among the differentially expressed genes identified by bulk proteomics across BN, MN, DH, DCIS, and IDC stages. (O) Boxplots showing THBS1 expression in bulk transcriptomics. (P) Boxplots showing MGP (left) and THBS1 (right) expression in bulk proteomics. Statistical analysis: Stage comparisons and statistical analyses were performed as described in Figure S2. All tests were performed using the t test for A, B, C, D, E, F, H, L. *P* values are presented to four significant digits; *P <* 0.0001 is indicated in scientific notation. BN, normal from benign patient; MN, normal from malignant patient; DH, intraductal hyperplasia; DCIS, ductal carcinoma in situ; IDC, invasive ductal carcinoma.

**Figure S4.**
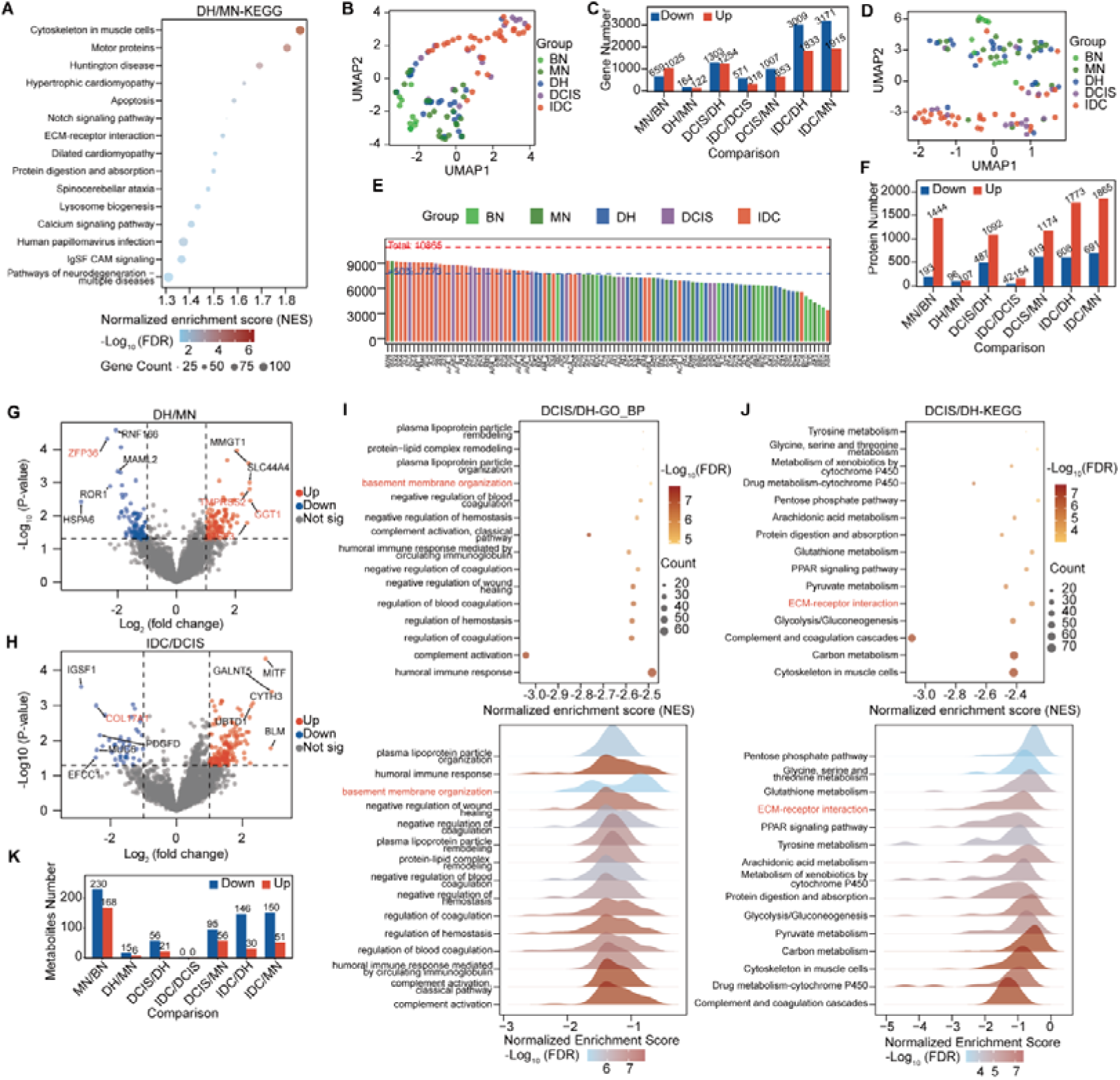
Multi-omics characterization of invasion-associated features in early breast cancer development, related to figure 4. (A) Bubble plot showing enriched KEGG pathways from GSEA for somatic mutations in DH compared to MN. Dot size indicates gene count, and color represents -log_10_(FDR) (blue to red, significant). (B) UMAP plot showing bulk transcriptomic sample distribution across BN, MN, DH, DCIS, and IDC stages. (C) Bar plot showing the number of differential transcriptomic genes across disease stages. (D) Bar plot showing protein identification numbers across disease stages in bulk proteomics. (E) UMAP plot showing bulk proteomic sample distribution across disease stages. (F) Bar plot showing the number of differential protein numbers across disease stages. (G) Volcano plots showing bulk proteomics changes during the MN to DH transition. (H) Volcano plots showing bulk proteomics changes during the DCIS to IDC transition. (I) Dot plot (upper) and ridge plot (lower) showing GO-BP GSEA results for DCIS v.s. DH. Significantly enriched terms (FDR < 0.05) are ranked by |NES| in descending order, with the top 15 terms shown. Terms are ordered by count in the dot plot (upper) and by NES in the ridge plot (lower). (J) Dot plot (upper) and ridge plot (lower) showing KEGG GSEA results for DCIS v.s. DH. Significantly enriched terms (FDR < 0.05) are ranked by |NES| in descending order, with the top 15 terms shown. Terms are ordered by count in the dot plot(upper) and by NES in the ridge plot (lower). Statistical analysis: Stage comparisons and statistical analyses were performed as described in Figure S2. For differential expression analysis in volcano plots of G and H, significance was defined as P value < 0.05 and |log_2_(fold change)| > log_2_(2). All enrichment analyses were performed using the clusterProfiler package. GSEA analysis was conducted using gseGO (BP) for I and gseKEGG for A and J, with a significance threshold of FDR < 0.05. BN, normal from benign patient; MN, normal from malignant patient; DH, intraductal hyperplasia; DCIS, ductal carcinoma in situ; IDC, invasive ductal carcinoma.

**Figure S5.**
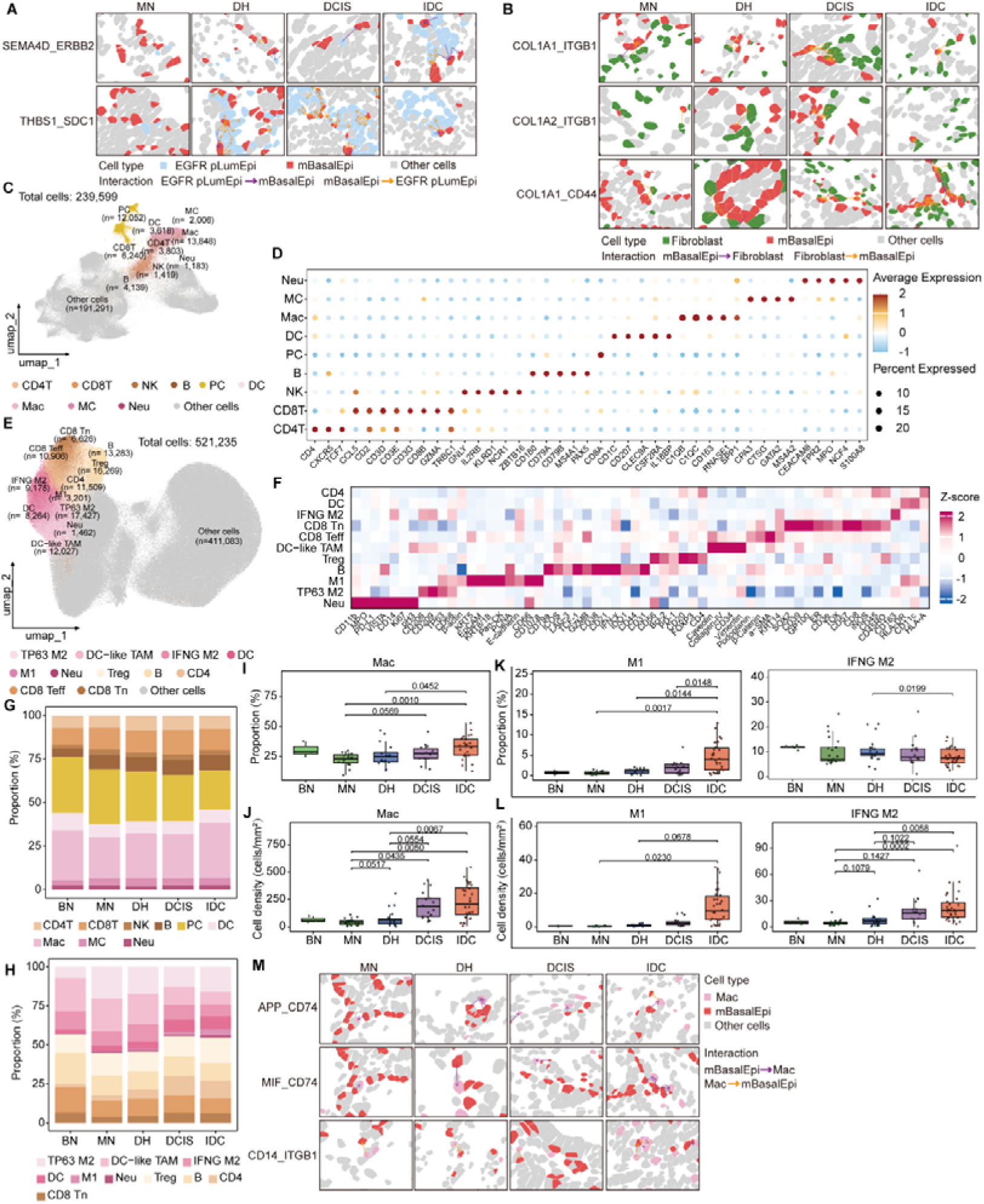
Immune cell subpopulation profiling and cell-cell interaction spatial validation during breast cancer progression, related to figure 5. (A) Representative spatial maps showing ligand-receptor interactions between mBasalEpi and EGFR pLumEpi (SEMA4D_ERBB2 and THBS1_SDC1) across MN, DH, DCIS and IDC stages in CosMx data; arrows indicate interaction direction. (B) Representative spatial maps showing ligand-receptor interactions between mBasalEpi and fibroblasts (COL1A1_ITGB1, COL1A2_ITGB1 and COL1A1_CD44) across MN, DH, DCIS and IDC stages in CosMx data; arrows indicate interaction direction. (C) UMAP projection of all 239,599 cells from CosMx data, colored by immune subclusters (other cells shown in gray). (D) Dot plot showing average expression level (color) and percentage of expressing cells (size) of marker genes for each immune subcluster in CosMx data. (E) UMAP projection of all 521,235 cells from CODEX data, colored by immune subclusters (other cells shown in gray) with a consistent color scheme as ST. (F) Heatmap of Z-score normalized marker protein expression across immune subclusters in CODEX data. (G) Stacked bar plot showing the proportion of immune subclusters across BN, MN, DH, DCIS and IDC stages in CosMx data. (H) Stacked bar plot showing the proportion of immune subclusters across disease stages in CODEX data. (I) Box-and-scatter plots depicting the proportion of Mac subcluster across disease stages in CosMx data. (J) Box-and-scatter plots showing cell density (cells/mm^2^) of Mac subcluster across disease stages in CosMx data. (K) Box-and-scatter plots depicting the proportion of M1 and IFNG M2 subclusters across disease stages in CODEX data. (L) Box-and-scatter plots showing cell density (cells/mm^2^) of M1 and IFNG M2 subclusters across disease stages in CODEX data. (M) Representative spatial maps showing ligand-receptor interactions between mBasalEpi and macrophages (APP_CD74, MIF_CD74 and CD14_ITGB1) across MN, DH, DCIS and IDC stages in CosMx ST data; arrows indicate interaction direction. Statistical analysis: Stage comparisons and statistical analyses were performed as described in Figure S2. All tests were performed using the t test for I-L. *P* values are presented to four significant digits; *P <* 0.0001 is indicated in scientific notation. BN, normal from benign patient; MN, normal from malignant patient; DH, intraductal hyperplasia; DCIS, ductal carcinoma in situ; IDC, invasive ductal carcinoma.

**Figure S6.**
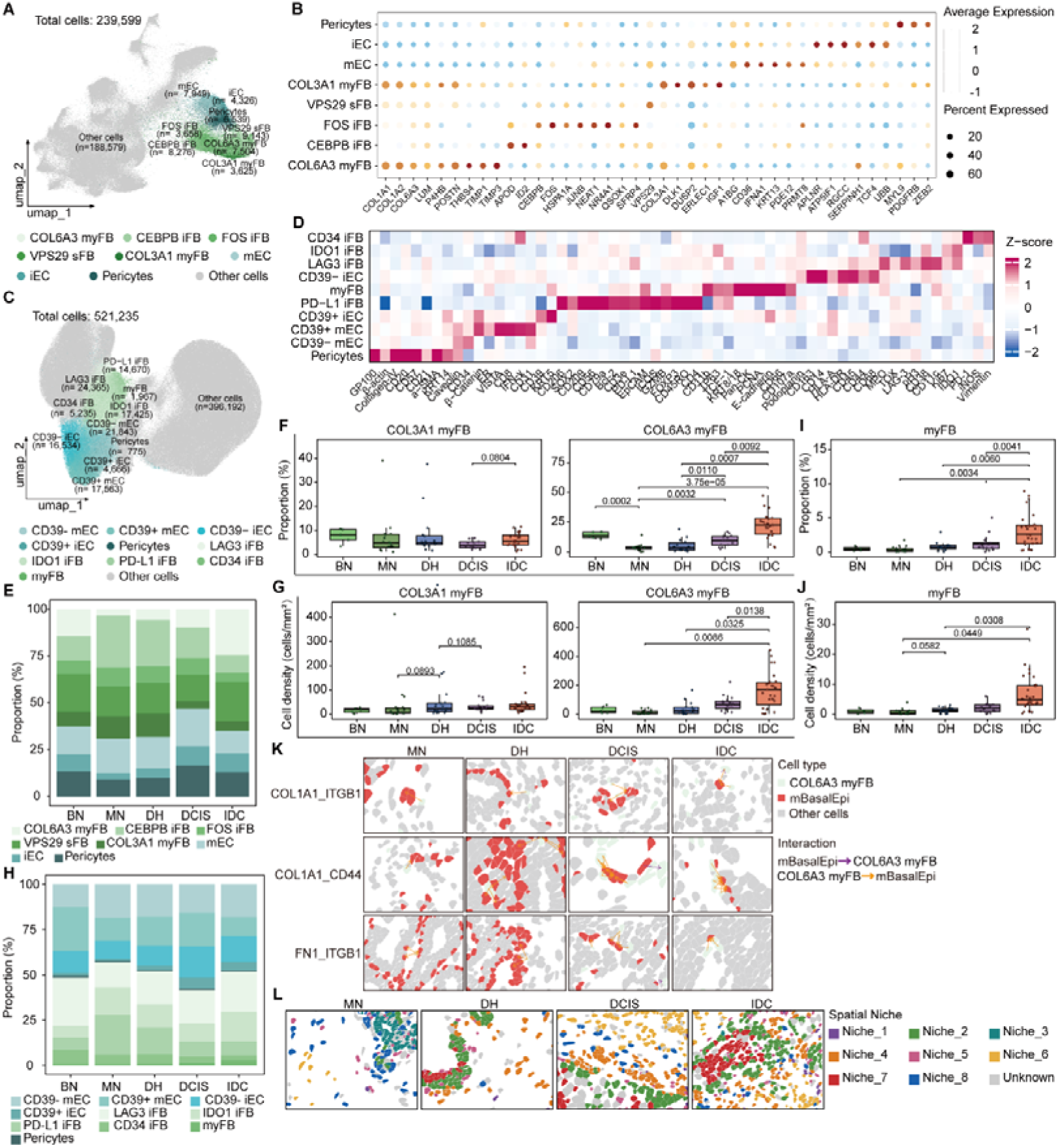
Stromal cell profiling, cell-cell interaction and spatial niche validation during breast cancer progression related to figure 5. (A) UMAP projection of all 239,599 cells from CosMx data, colored by stromal cell (other cells shown in gray). (B) Dot plot showing average expression level (color) and percentage of expressing cells (size) of marker genes for each stromal cell subcluster in CosMx data. (C) UMAP projection of all 521,235 cells from CODEX data, colored by stromal cell subclusters (other cells shown in gray) with a consistent color scheme as CosMx. (D) Heatmap of Z-score normalized marker protein expression across stromal cell subclusters in CODEX data. (E) Stacked bar plot showing the proportion of stromal cell subclusters across BN, MN, DH, DCIS and IDC stages in CosMx data. (F) Box-and-scatter plots depicting the proportion of COL3A1 myFB and COL6A3 myFB subclusters across disease stages in CosMx data. (G) Box-and-scatter plots showing cell density (cells/mm^2^) of COL3A1 myFB and COL6A3 myFB subclusters across disease stages in CosMx data. (H) Stacked bar plot showing the proportion of fibroblast subclusters across disease stages in CODEX data. (I) Box-and-scatter plots depicting the proportion of myFB subcluster across disease stages in CODEX data. (J) Box-and-scatter plots showing cell density (cells/mm^2^) of myFB subcluster across disease stages in CODEX data. (K) Representative spatial maps showing ligand-receptor interactions between mBasalEpi and COL6A3 myFB (COL1A1_ITGB1, COL1A1_CD44 and FN1_ITGB1) across MN, DH, DCIS and IDC stages in CosMx data; arrows indicate interaction direction. (L) Representative spatial distribution maps of cellular niches across MN, DH, DCIS and IDC stages in CosMx data. Statistical analysis: Stage comparisons and statistical analyses were performed as described in Figure S2. All tests were performed using the t test for F, G, I, J. *P* values are presented to four significant digits; *P <* 0.0001 is indicated in scientific notation. BN, normal from benign patient; MN, normal from malignant patient; DH, intraductal hyperplasia; DCIS, ductal carcinoma in situ; IDC, invasive ductal carcinoma.

**Figure S7.**
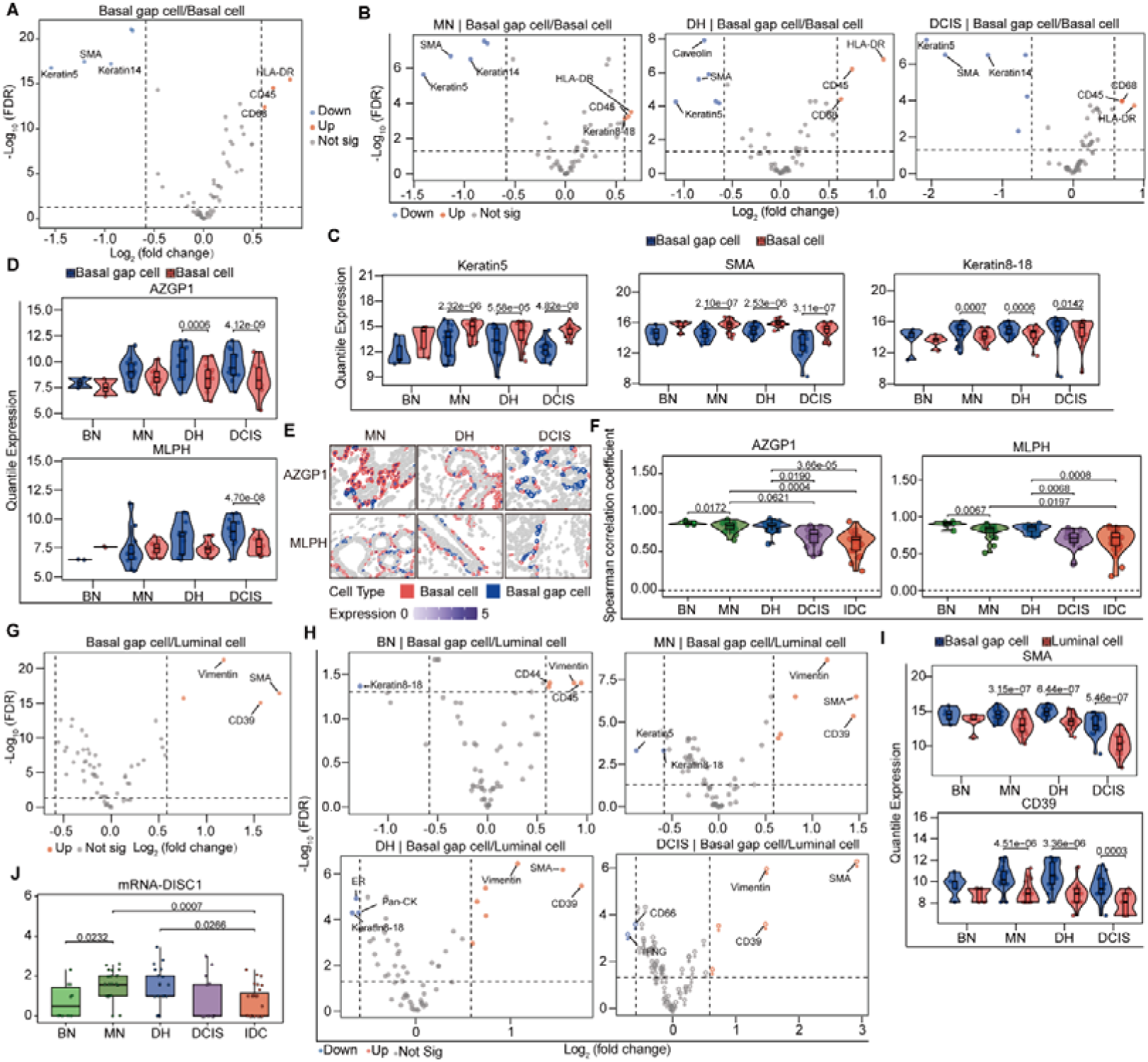
Luminal leader cells breach the basal cell layer to generate gaps, related to figure 6. (A) Volcano plot showing differentially expressed proteins (DEPs) between basal gap cells and basal cells across all disease stages in CODEX data. (B) Volcano plots showing DEPs between basal gap cells and basal cells stratified by disease stage (MN, DH, DCIS) in CODEX data. (C) Violin-box-and-scatter plots depicting the expression levels of representative DEPs (Keratin5, SMA, Keratin8-18) in basal gap cells and basal cells across disease stages in CODEX data; *P* values represent FDR from differential analysis. (D) Violin-box-and-scatter plots depicting the expression levels of representative DEGs (AZGP1 and MLPH) in basal gap cells and basal cells across disease stages in CosMx data; *P* values represent FDR from differential analysis. (E) Representative spatial expression maps of the above DEGs in basal gap cells and basal cells across disease stages, with cell type indicated by border color and gene expression level represented by fill intensity. (F) Violin-box-and-scatter plots depicting the spearman correlation coefficient of protein AZGP1 and MLPH in SMA+ cells across disease stages. (G) Volcano plot of DEPs between basal gap cells and luminal cells across all disease stages in CODEX data. (H) Volcano plots of DEPs between basal gap cells and luminal cells stratified by disease stage (BN, MN, DH, DCIS) in CODEX data. (I) Violin-box-and-scatter plots depicting the expression levels of representative DEPs (SMA, CD39) in basal gap cells and luminal cells across disease stages in CODEX data; *P* values represent FDR from differential analysis. (J) Boxplots showing DISC1 mRNA expression levels across disease stages in bulk transcriptomic. Statistical analysis: Stage comparisons and statistical analyses were performed as described in Figure S2. For differential expression analysis in volcano plots (A, B, G, H), significance was defined as FDR < 0.05 and |log_2_(fold change)| > log_2_(1.5). All tests were performed using the limma for A, B, C, D, G, H, I and t test for F, J. *P* values are presented to four significant digits; *P <* 0.0001 is indicated in scientific notation. BN, normal from benign patient; MN, normal from malignant patient; DH, intraductal hyperplasia; DCIS, ductal carcinoma in situ; IDC, invasive ductal carcinoma.

